# Subcortical Atlas of the Rhesus Macaque (SARM) for Magnetic Resonance Imaging

**DOI:** 10.1101/2020.09.16.300053

**Authors:** Renée Hartig, Daniel Glen, Benjamin Jung, Nikos K. Logothetis, George Paxinos, Eduardo A. Garza-Villarreal, Adam Messinger, Henry C. Evrard

**Author notes:** Corresponding author: Henry C. Evrard, Nathan S. Kline Institute for Psychiatric Research, 140 Old Orangeburg Road, Orangeburg, New York 10962, USA.

## Abstract

Digitized neuroanatomical atlases are crucial for localizing brain structures and analyzing functional networks identified by magnetic resonance imaging (MRI). To aid in MRI data analysis, we have created a comprehensive parcellation of the rhesus macaque subcortex using a high-resolution *ex vivo* structural imaging scan. The structural scan and its parcellation were warped to the updated <u>NIMH Macaque Template</u> (NMT v2), an *in vivo* population template, where the parcellation was refined to produce the Subcortical Atlas of the Rhesus Macaque (SARM). The subcortical parcellation and nomenclature reflect those of the 4^th^ edition of the Rhesus Monkey Brain in Stereotaxic Coordinates (RMBSC4; Paxinos et al., in preparation). The SARM features six parcellation levels, arranged hierarchically from fine regions-of-interest (ROIs) to broader composite regions, suited for fMRI studies. As a test, we ran a functional localizer for the dorsal lateral geniculate (DLG) nucleus in three macaques and found significant fMRI activation in this atlas region. The SARM has been made openly available to the neuroimaging community and can easily be used with common MR data processing software, such as AFNI, where the atlas can be embedded into the software alongside cortical macaque atlases.

**Highlights:**

- We present the Subcortical Atlas of the Rhesus Macaque (SARM).
- SARM provides a neuroanatomical reference frame for neuroimaging analysis.
- The entire subcortex is mapped, including the thalamus, basal ganglia, and brainstem.
- ROIs are grouped hierarchically, making SARM useful at multiple spatial resolutions.
- SARM is in the NMT v2 template space and complements the CHARM atlas for the cortex.

## 1. Introduction

As functional magnetic resonance imaging (fMRI) continues to advance spatiotemporal resolution limits, there is a growing opportunity for researchers to examine subcortical regions and their involvement in cortico-subcortical networks. These smaller subcortical regions have, however, largely been absent from digitized atlases applicable to MRI research with non-human primates (NHPs). In contrast to human research, where several subcortical atlases exist, NHP researchers typically have to employ workarounds and parcellate individual regions of interest (ROIs) themselves. To address this void, we present the Subcortical Atlas of the Rhesus Macaque (SARM), a digital subcortical atlas offering a standardized parcellation for ROI and network analyses.

The development of the SARM is timely. While previously used in only a few primate research centers, fMRI is now being employed in many NHP laboratories (Milham et al., 2018). The use of contrast agents, improved sequences, and high-field magnets is increasing the signal-to-noise ratio and spatial resolution into the realm where subcortical activations can be reproducibly detected (e.g., Baker et al., 2006; Ortiz-Rios et al., 2015; Quan et al., 2020). Technological improvements in data collection methods have also resulted in greater potential for employing fMRI concurrently with subcortical electrical microstimulation (Logothetis et al., 2010; Arsenault & Vanduffel, 2019; Murris, Arsenault & Vanduffel, 2020), optogenetics (Nassi et al., 2015; Klein et al., 2016; Stauffer et al., 2016), or electrophysiological recordings (Logothetis et al., 2012), all the while capturing the mesoscopic and systems-level effects (see also, Klink et al., this issue). Such studies require fine-grain delineations of the subcortex to aid both in planning stereotaxic implantations and interpreting local signal modulation. Finally, while NHP fMRI still typically relies on two or three subjects, there is a growing interest in using larger groups and applying group analyses (e.g. Fox et al., 2018). The advent of multi-center data sharing (Milham et al., 2018) also allows for the possibility of larger sample sizes, and clearly calls for group-level analyses performed on data aligned to a population brain template with standardized atlases (Milham et al., 2020; Jung et al., this issue).

Previous NHP studies examining subcortical activity have created their own individual masks covering regions known to include specific brainstem nuclei. For example, Logothetis et al. (2012) manually segmented 25 subcortical ROIs for each of their five subjects separately. Noonan et al. (2014) masked the area between the medulla and midbrain for localizing activity from serotonergic nuclei with 0.5 mm spatial resolution. Murris et al. (2020) registered their functional data to the D99 macaque template (Reveley et al., 2017) and then added ROIs for the ventral tegmental area (VTA) and accumbens nucleus (Acb), which are absent in the D99 atlas. While creating individual masks is one strategy to study regional fMRI activity, precise delineation of structural boundaries requires not only high-quality structural scans but a great deal of labor and anatomical expertise. Furthermore, for comparisons across individuals or group-level analysis, single-subject scans and regional masks must then be nonlinearly registered to a common reference template. While warping of fMRI data to a standard space can be successful for fairly large subcortical parcellations (e.g., the hippocampus, amygdala, and distinguishable midbrain regions; Fox et al., 2015), nonlinear registration of smaller subcortical structures can be a delicate step in the data processing pipeline. The SARM allows for varying alignment and resolution limits, providing hierarchically arranged groupings of regions that are suited for different purposes, including functional neuroimaging studies.

In comparison to human MRI brain atlases (e.g., Accolla et al., 2014; Pipitone et al., 2014; Ewert et al., 2017; Pauli et al., 2018), and despite the existence of NHP paper atlases including an exhaustive mapping of the subcortex (Paxinos et al., 2009; Martin & Bowden, 2000), limited efforts have been made to digitize the macaque subcortex parcellations. Previous attempts to digitize subcortical parcellations from printed macaque atlases have provided some segmentation of the subcortex. For example, the Saleem and Logothetis (2012) atlas was digitized by alignment to a high-resolution MRI of an *ex vivo* surrogate (Reveley et al. 2017). However, this digital D99 atlas includes only some subcortical structures (e.g., hippocampus, amygdala, striatum, and claustrum). Likewise, the parcellation of post-mortem macaque brains by Calabrese et al. (2015) contains a detailed segmentation of most telencephalic and diencephalic brain nuclei (Paxinos et al., 2009) but little to no parcellation of the brainstem. The NeuroMaps macaque atlas covers the whole brain and is presented on an *ex vivo* juvenile rhesus macaque brain (Bakker, Tiesinga & Kötter, 2015; Rohlfing et al., 2012). The NeuroMaps segmentation was later refined on the INIA19 adult population *in vivo* symmetric template (Rohlfing et al., 2012). Rohlfing and colleagues noted, however, that their segmentation of the basal forebrain, hypothalamus and amygdala is incomplete and that the internal segmentation of the thalamus, midbrain and hindbrain may not be reliable. Although a more detailed digital subcortical map is needed, segmenting all the minute cytoarchitectonic subnuclei that can be appreciated under the microscope would be of little value at fMRI resolution. Using an updated version of the Rhesus Monkey Brain in Stereotaxic Coordinates (Paxinos et al., 2009; Paxinos et al., in preparation) for guidance, the SARM addresses the need for a more comprehensive subcortical segmentation while attempting to strike a practical balance between anatomical details and the constraints imposed by the lower anatomical resolution of MRI.

While *ex vivo* scans, such as the D99 surrogate or the Calabrese *ex vivo* population template (Calabrese et al., 2015), can provide great detail because they are not impacted by animal movement or physiological noise, *in vivo* templates better reflect the living brain’s configuration (e.g., with regard to size, ventricle shape, and the presence of cerebrospinal fluid). Atlases drawn on a single subject template can precisely reflect the particular anatomy of that subject but may not be morphologically representative of the species due to large inter-individual variability. The SARM was fit to version 2 of the NIMH Macaque Template (NMT v2), a high-resolution population template based on *in vivo* scans collected at high field strength (4.7 T) from a large cohort (N=31) of adult rhesus monkeys (Seidlitz et al, 2018; Jung et al., this issue). The NMT v2 compares favorably to the INIA19 template in terms of resolution (0.25 mm vs. 0.50 mm isotropic), allowing for finer parcellation of subcortical structures, and is already home to the Cortical Hierarchy Atlas of the Rhesus Macaque (CHARM; Jung et al., this issue). Because average population templates are representative, most individuals will require relatively little distortion to be aligned to such a template as compared to an *ex vivo* or individual scan (Kochunov et al., 2001; Molfese et al., 2015; Feng et al., 2017). This, in effect, minimizes alignment errors, which are of particular importance for small subcortical nuclei.

To create the SARM, we relied on the high resolution and precision of an *ex vivo* single-subject structural scan and previously obtained histological material to draw the primary structures. These regions were then warped to the symmetric version of the NMT v2, where they were manually refined to reflect the representative anatomy of the population template. The SARM parcellation was then hierarchically grouped into larger composite structures to create region of interest (ROI) clusters suitable for (f)MRI analysis. The SARM is available on various online platforms (<u>PRIME-RE</u>, <u>Zenodo</u>, and <u>AFNI</u>), where it is being continuously improved and further delineated.

## 2. Materials & Methods

### 2.1 Atlas Preparation

#### 2.1.1 Ex Vivo Anatomical Sample

A whole-brain *ex vivo* sample from one adult female rhesus macaque (G12; *Macaca mulatta*; ∼8 kg) was used as a single-subject anatomical template to parcellate the subcortex. This subject was part of an anatomical study approved by the local authorities and in full compliance with the European Parliament and Council Directive 2010/63/EU. The subject was not involved in any invasive procedures and never underwent intracerebral surgery. After transcardial fixation with 4% formalin (Evrard et al., 2012), the brain was placed into a jar of agar and positioned upright in a horizontal 7T Bruker BioSpec scanner, with the brain oriented parallel to the scanner (dorsal side positioned upward) (Bruker BioSpin, Ettlingen, Germany). The entire brain was scanned using a high-resolution fast low-angle shot (FLASH) sequence (voxel dimensions: 0.15×0.15×1.0 mm; flip angle: 50°; TR/TE: 2500/9 msec; field-of-view (FOV): 70×52 mm; matrix size: 468×346; 78 coronal slices).

#### 2.1.2 Segmentation in Individual (G12) Space

Subcortical ROIs were manually drawn by author HCE onto coronal slices of the G12 high-resolution *ex vivo* anatomical scan using the Amira software (Amira 6.0.1; FEI). The fine spatial resolution of the contrast variation in the slices enabled recognizing and mapping discrete anatomical regions identified in corresponding histological sections from the 4^th^ edition of the Rhesus Monkey Brain in Stereotaxic Coordinates (RMBSC4; Paxinos et al., in preparation). The order of the figures in this upcoming edition does not differ from the 2^nd^ edition; thus readers can still refer to the printed second edition of RMBSC (Paxinos et al., 2009) when references to specific figures are made in the text below. These reference sections previously underwent Nissl and AChE staining (Paxinos et al., 2009), and were recently scanned using a slide-scanner microscope (AxioScan; Zeiss) for further examination (Paxinos et al., in preparation). The ROIs were drawn while examining all three stereotaxic planes to reduce inconsistencies in delineation across slices. Regions defined in RMBSC4 that were too small and not clearly discernible from changes in contrast in the G12 scan were grouped together in larger ROIs, as detailed in the Results (Section 3.1). Additional resources included prior architectonic parcellations of the hypothalamus (Saper et al., 2012), thalamus (Olszewki, 1952; Calzavara et al., 2005; Evrard and Craig, 2008; Mai and Forutan, 2012), amygdala (Amaral et al., 1992; Stefanacci et al., 2000), and basal ganglia (Haber et al., 2012).

##### 2.1.3 Nonlinear Registration

The single-subject (G12) *ex vivo* structural scan and subcortical segmentation were nonlinearly registered to the symmetric NMT v2 full-head anatomical template for rhesus macaques. The NMT v2 template (Jung et al., this issue) is in stereotaxic orientation (Horsley and Clarke, 1908; also referred to as the Frankfurt Zero plan). The subcortical segmentation was refined on a single hemisphere (the left) of the NMT v2 and mirrored onto the opposite hemisphere in order to assure that the resulting parcellation has left and right ROIs of equal size.

To coregister the G12 template and atlas to the NMT v2, the NIFTI images were first converted to MINC format (http://www.bic.mni.mcgill.ca/ServicesSoftware/MINC) and the origin of the spatial coordinates was adjusted to correspond to the intersection of the midsagittal section and the interaural line (i.e., ear bar zero, EBZ). Then, we used *volmash* and *volflip* (<u>MINC widgets</u>) to reorient the images to the NMT v2. The G12 template was then converted back to NIFTI. Using Advanced Normalization Tools (ANTs; version 2.3.1.dev159-gea5a7; Avants et al., 2014), we made a negative image of the G12 so its contrast would be similar to the T1-weighted NMT v2 template. The G12 template showed air bubble-induced artifacts around the left lateral ventricle that affected registration. To correct these artifacts and improve registration, we manually traced each artifact to the underlying tissue (namely, the putamen) and matched it with the tissue’s intensity. This new volume was then corrected for N4 Bias Field artifacts (Tustison et al., 2010). The ANTs registration pipeline was optimized using an in-house script that employed a custom mask of the subcortex for some of the registration steps. After computing the G12 to NMT v2 template registration, we used *antsApplyTransformation* to nonlinearly coregister the subcortical parcellation to the NMT v2 with Generic Label interpolation.

##### 2.1.4 Refinement of ROIs in the NMT v2 Template

The resulting atlas regions suffered from some irregularities stemming from the limitations of the original anisotropic voxels (high resolution within the coronal plane, but coarser resolution across planes) and from the interpolation methods associated with the nonlinear warp of the ROI labels. Therefore, we followed the ANTs-based alignment pipeline with a procedure to spatially regularize regions using AFNI commands. The regions were processed with a modal smoothing technique that replaces each voxel with the most common label in a 1- or 2-voxel spherical neighborhood around every voxel. A select list of thin or small regions were smoothed using the 1 voxel mode, and all other regions were smoothed using the 2 voxel mode. The data were masked by the CSF and blood vessel segmentations from NMT v2. Each ROI was automatically further refined by examining the distribution of voxel intensities in NMT v2. For each ROI, we sampled voxel intensities of NMT v2 in that ROI, and voxels farther than three standard deviations away from the mean intensity (potentially indicating encroachment of the ROI into a different tissue class) were compared with eight neighboring voxels and reassigned to the label of the voxel with the most similar intensity. This outlier detection was performed across ten iterations. The quality of the alignment between the transformed G12 and the NMT v2 template was assessed by viewing the former on the outline of the latter using @chauffeur_afni. Finally, the atlas was assessed for discontinuities, and discontinuous clusters smaller than five percent of the size of the largest portion of the ROI were replaced with labels from neighboring voxels. With the atlas regions now transformed to the NMT v2 symmetric template space, the regions were manually adjusted, again in Amira by author HCE, with reviewing by authors HCE and GP, to reflect the anatomical transitions evident in this population template. Before exporting from AMIRA, a Gaussian smoothing (2×2×2 pixel filter mask) was applied across the 3D volume using the “Smooth Labels” function. Following AMIRA export, the SARM regions were modally smoothed with a 1.8 voxel radius and discontinuous clusters smaller than five percent of the size of the largest portion of the ROI were again replaced with labels from neighboring voxels. At each step, volume changes of each ROI were tracked to prevent large, unintended changes to the ROIs.

### 2.2 Subcortical Naming Hierarchy

#### 2.2.1 Hierarchical Grouping

To create ROIs of varying spatial resolution, the neighboring regions in the primary parcellation were iteratively grouped to form a hierarchy of subcortical structures across six levels that describes progressively larger and more general anatomical regions. This hierarchy forms the SARM. The finest level of the SARM hierarchy (level 6) individually itemizes each of our manually drawn ROIs, which were defined on the basis of the RMBSC4, as described above. Composite regions in levels 1-5 were successively built from smaller adjacent areas in the next finer level. While the brain can most broadly be subdivided into the forebrain, midbrain, and hindbrain, level 1 begins with the developmental and embryological sub-divisions of the subcortex, namely the tel-, di-, mes-, met-, and myel-encephalon. The SARM levels 2-4 consist of ROIs of sufficient size to accommodate functional imaging voxels that are typically 1.25-1.50 mm on a side, whereas levels 5-6 ROIs may benefit from the higher resolution of structural imaging. Levels 5 and 6 of the SARM were left largely similar to allow for potential future delineation of SARM regions. In most cases, we grouped the ROIs based on their developmental and/or functional relationships (Mai and Paxinos, 2012; Puelles et al., 2013; see also Calabrese et al., 2015 for a similar approach), with the condition that these ROIs had to be spatially contiguous. In other cases, in particular at coarser levels, ROIs had to be grouped solely based on their spatial proximity.

Independent of their hierarchical classification, all ROIs were classified as being primarily subcortical gray matter or white matter. In select instances, ROIs composed primarily of white matter or other tissue types were included in larger composite ROIs to make them whole (e.g. the internal capsule was included in the striatum to bridge the caudate and putamen) and because sparse cell bodies within such white matter regions can lead to their functional activation.

#### 2.2.2 Nomenclature

Each ROI and group of ROIs has a unique full name and abbreviation. At levels 5 and 6, the names and abbreviations of the ROIs typically match those defined in the RMBSC4, with some exceptions (see Results) to accommodate the most commonly used naming convention in NHP fMRI. At levels 2 to 4, the names and abbreviations of the groups of ROIs reflect either a common developmental origin (e.g., pallial vs. subpallial amygdala; Puelles et al., 2013), a classical neuroanatomical grouping (e.g., basal ganglia; Mai and Paxinos, 2012) or a spatial proximity (e.g., dorsal vs. ventral mesencephalon).

AFNI allows for flexible indexing of ROIs by either index number, abbreviation, or the full name of the ROI. To prevent conflicts between index numbers and names, SARM abbreviations do not start with a number (e.g., the abducens nucleus is abbreviated 6N in RMBSC4 but N6 in SARM). In addition, to maximize compatibility with scripts and programs, abbreviations do not include special characters, and full names use underscores in place of spaces. A full list of the current SARM regions is provided in **Supplementary Table 1**. A spreadsheet of the hierarchy and full list of SARM structures is also available for download with the NMT v2 package.

### 2.3 Functional Localizer

To illustrate the usefulness of this atlas within the context of fMRI data analysis, a functional localizer for the dorsal lateral geniculate nucleus (DLG) (also referred to as the lateral geniculate nucleus, LGN) was included, from a larger experimental program, with three adult rhesus macaque*s* (*Macaca mulatta*; 1 female; average weight: 10.11 kg). Experiments were conducted following a previously described opiate-based anesthesia protocol (Logothetis et al., 2010). Animals were treated according to the guidelines of the European Parliament and Council Directive 2010/63/EU on the protection of animals used for experimental and other scientific purposes. Experimental protocols were approved by the local German authorities.

#### 2.3.1 Image Acquisition

Neuroimaging data were acquired using a vertical 7 Tesla NMR scanner (Bruker, Billerica, MA, U.S.A.) and Paravision software (version 5). fMRI data were acquired with a quadrature coil and double-shot gradient-echo echo planar imaging (GE-EPI; voxel dimensions: 0.75×0.75×2.0 mm; flip angle: 53°; TR/TE: 2000/19 msec; FOV: 96×96 mm; matrix size: 128×128; 20 axial slices). Slice volumes were acquired contiguously. During each experiment, a T2-weighted rapid acquisition with relaxation enhancement (RARE) scan was collected to image the native structural space (RARE factor: 8; voxel dimensions: 0.375×0.375×1.0 mm; flip angle: 180°; TR/TE: 6500-8500/16 msec; FOV: 96×96 mm; matrix size: 256×256; 40 axial slices). Acquired data were converted offline from Bruker file format to 4D NIFTI files using the Unix-based *<u>pvconv</u>*.

#### 2.3.2 Stimulus

A flickering checkerboard stimulus was visually presented (Logothetis et al., 1999) during a 10 minute GE-EPI scan, consisting of 300 volumes. The stimulus was presented for 4 sec preceded by an 8 sec OFF period, and followed by a longer 18 sec OFF period, allowing return of the blood-oxygen-level-dependent (BOLD) signal to baseline. Two sessions per subject were collected and analyzed using two common software packages (SPM & AFNI) to validate the application of SARM for studying subcortical activity across different processing pipelines.

#### 2.3.3 SPM-Based Image Analysis

Functional data were realigned using SPM12 (Statistical Parametric Mapping; Wellcome Department of Imaging Neuroscience, London, UK) to obtain six rigid-body transformation parameters and then aligned to each subject’s native anatomical (RARE) scan. Each subject’s RARE was subsequently translated to the NMT v2 space, and this linear transformation was applied to all relevant functional scans. Data were nonlinearly aligned using SPM-based Dartels, a diffeomorphic warping algorithm (Ashburner, 2007), which relies on tissue class identification and segmentation. The resulting deformation matrix was applied to each individual’s RARE and fMRI images. At each step, the spatial alignment was checked by direct visual examination. The EPIs were smoothed (2 mm FWHM Gaussian) and the fMRI data were estimated using a General Linear Model (GLM), which included as regressors the rigid-body transformation parameters, in the event-related responses correlated with the visual stimulus presentation (*B*_1_) and the baseline activity (*B*_o_). The fMRI data were averaged across sessions for each subject, and significant activations were assessed with a T-contrast (*p* < 0.05, FDR-corrected).

#### 2.3.4 AFNI-Based Image Analysis

Using AFNI (Cox, 1996), functional data were processed by first computing the alignment of each subject’s T2 structural (RARE) scan to the NMT v2 template using the @animal_warper pipeline (Jung et al., this issue). To address the contrast (e.g., of CSF, GM and WM) profile inversion between NMT v2 and the functional localizer datasets, we used an alignment method that identifies local negative correlations to minimize the cost function, the Local Pearson Correlation (lpc; Saad, 2009). This cost function was used for both affine and nonlinear alignment. The alignment was assessed using AFNI visualization tools. The affine and nonlinear transformations and the skull-stripped dataset served as the input to the functional processing performed by afni_proc.py. This processing used typical options for motion correction, alignment of the functional data to the individual’s T2 anatomical dataset, and modal smoothing (by 1 voxel), followed by a per-voxel mean scaling. The normalized functional data were interpolated to an isotropic voxel resolution of 1.25 mm^3^. The functional paradigm was modeled using a BLOCK hemodynamic response function model, stimuli convolved with a 4-sec duration boxcar function and normalized to unit size.

### 2.4 Data Accessibility and Availability

The SARM and NMT v2 files are provided in NifTI and GifTI file format for compatibility with most neuroimaging programs. This package also includes: the original G12 dataset with ROI drawings in their original space and the full list of SARM ROIs, abbreviations, and grouping levels. For data transformation and analysis, relevant scripts are also provided. All resources described are currently openly available or will be made available in the near future through the PRIME-Resource Exchange (https://prime-re.github.io/) (Messinger et al., this issue), Zenodo (https://zenodo.org/record/4026520#.X10X95P0nlw), and can be downloaded along with the NMT v2 from the AFNI website (https://afni.nimh.nih.gov/pub/dist/doc/htmldoc/nonhuman/macaque_tempatl/atlas_sarm.html) or using the AFNI command @Install_NMT.

## 3. Results

### 3.1 Subcortical ROI Segmentation and Hierarchical Grouping

This first version of the SARM (SARM v1) contains 206 primary subcortical ROIs. These ROIs were first drawn on the G12 high-resolution scan and then nonlinearly aligned to the NMT v2 population-averaged symmetrical template. Most of these ROIs were anatomically identifiable in G12 and, to some extent, in NMT v2, based on local signal contrast variations. Regions likely to be relevant for MRI analyses, but not readily identifiable in either scan, were delineated based on their most likely topological localization and neighborhood relationships, using RMBSC4 as the principal reference (Paxinos et al., 2009; Paxinos et al., *in preparation*). Individual ROIs represent either a single homogeneous anatomical entity, as defined in RMBSC4, or a collection of smaller cytoarchitectonic entities that could not be distinguished from one another due to a lack of contrast differentiation. Beyond the definition of the manually drawn primary ROIs, we created ROIs of progressively larger size by successively aggregating primary ROIs across six hierarchical levels. These six levels can accommodate structural and functional neuroimaging datasets of various spatial resolutions and analyses at different degrees of anatomical detail. The following sections report, successively, the alignment of G12 to NMT v2 (Section 3.1.1), general observations on the hierarchical grouping of the ROIs (Section 3.1.2), and, finally, an overview of the definition of the individual ROIs and their hierarchical groups (Section 3.1.3).

#### 3.1.1 Alignment of G12 to NMT v2

Figure 1 portrays the G12 *ex vivo* anatomical MRI in stereotaxic space, corresponding to the NMT v2 template at 5 representative coronal sections, where the nonlinear alignment computed using ANTs was overlaid onto the edge contours of the NMT v2 template.

**Figure 1.**
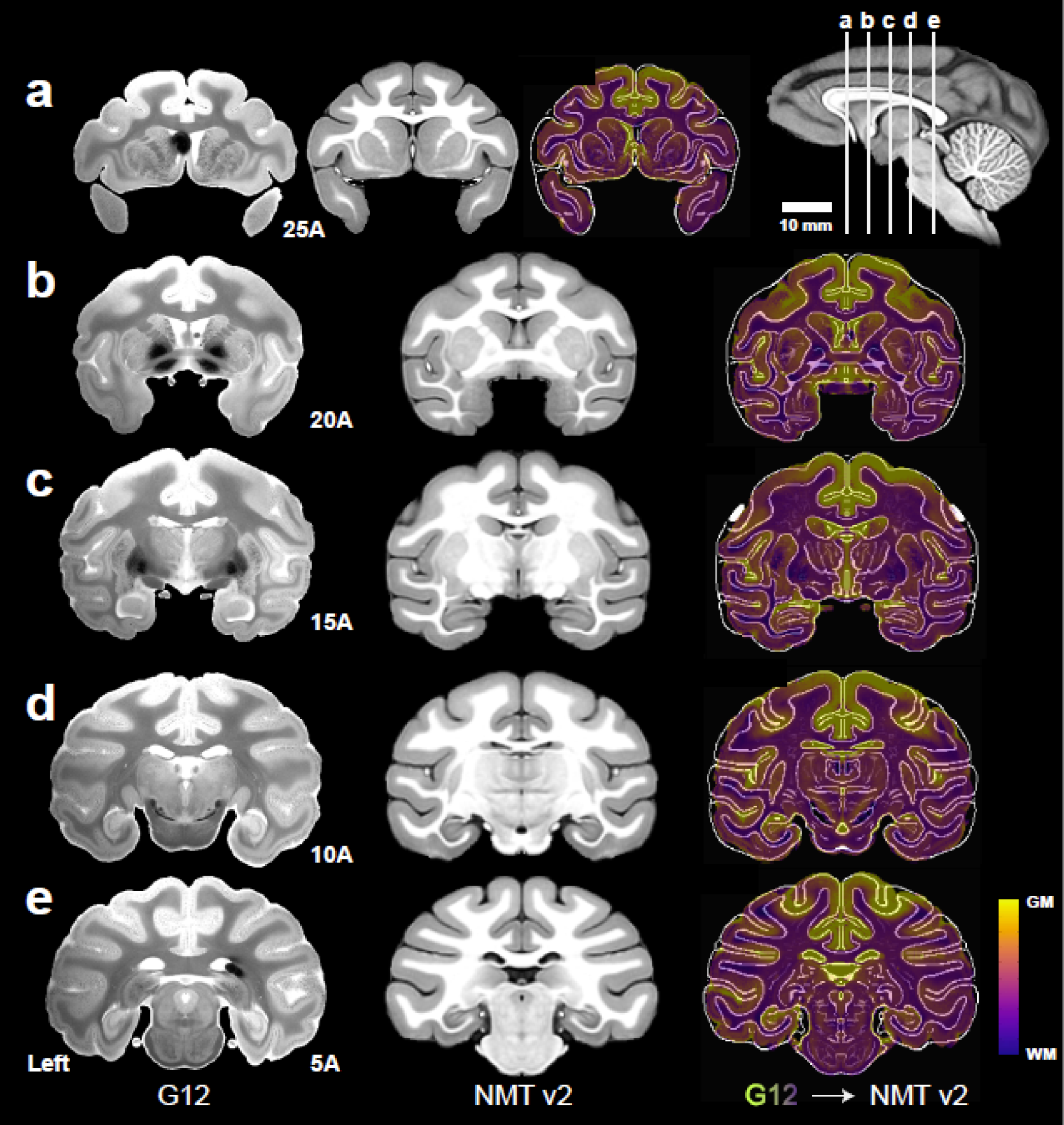
Alignment of subject G12 to the symmetric NMT v2. Panels (**a-e**) depict coronal slices through the G12 anatomical scan (left), the symmetric NMT v2 (middle) in stereotaxic space, and the nonlinear registration of the G12 to the symmetric template (right). Slice positions are in mm anterior to the origin (EBZ; ear bar zero) and are depicted on the midsagittal NMT v2 cross-section (upper right). Parameters for the ANTs registration pipeline were customized to prioritize alignment of subcortical regions. Color (‘plasma’) shows the warped G12 tissue intensities superimposed on the salient edges of the NMT v2. The darkest purple represents white matter (WM), whereas lighter purple and greens represent gray matter (GM). Left hemisphere depicted on the left side.

The G12 scan and its subcortical labels were resampled to match the NMT v2 resolution (0.25 mm^3^ isotropic), and the out-of-plane detail in the G12 was interpolated to match the NMT v2 resolution. Differences in morphology due to their preparations were noted (i.e., sulcal positioning, ventricle size and *ex vivo* fixation effects, as well as the presence of artifacts like air bubbles), because these may result in nonlinear registration errors or require repositioning greater than what is allowed by the nonlinear registration algorithm (cost function tradeoff).

The subcortical labels aligned to the NMT v2 exhibited some small irregularities along the region edges. These were mitigated with modal smoothing, clustering and outlier detection to make for more natural, locally consistent regions. At this stage, the regions underwent manual correction, followed by additional post-processing (see Section 2.1.4). The 3D consistency of the completed ROIs was verified by visualizing the surface of each region using AMIRA and AFNI’s surface viewer SUMA (Saad et al., 2004) (Fig. 2). The primary subcortical parcellation that forms the finest hierarchical level of the SARM (level 6; see Section 3.1.2) is shown in Figure 3.

**Figure 2.**
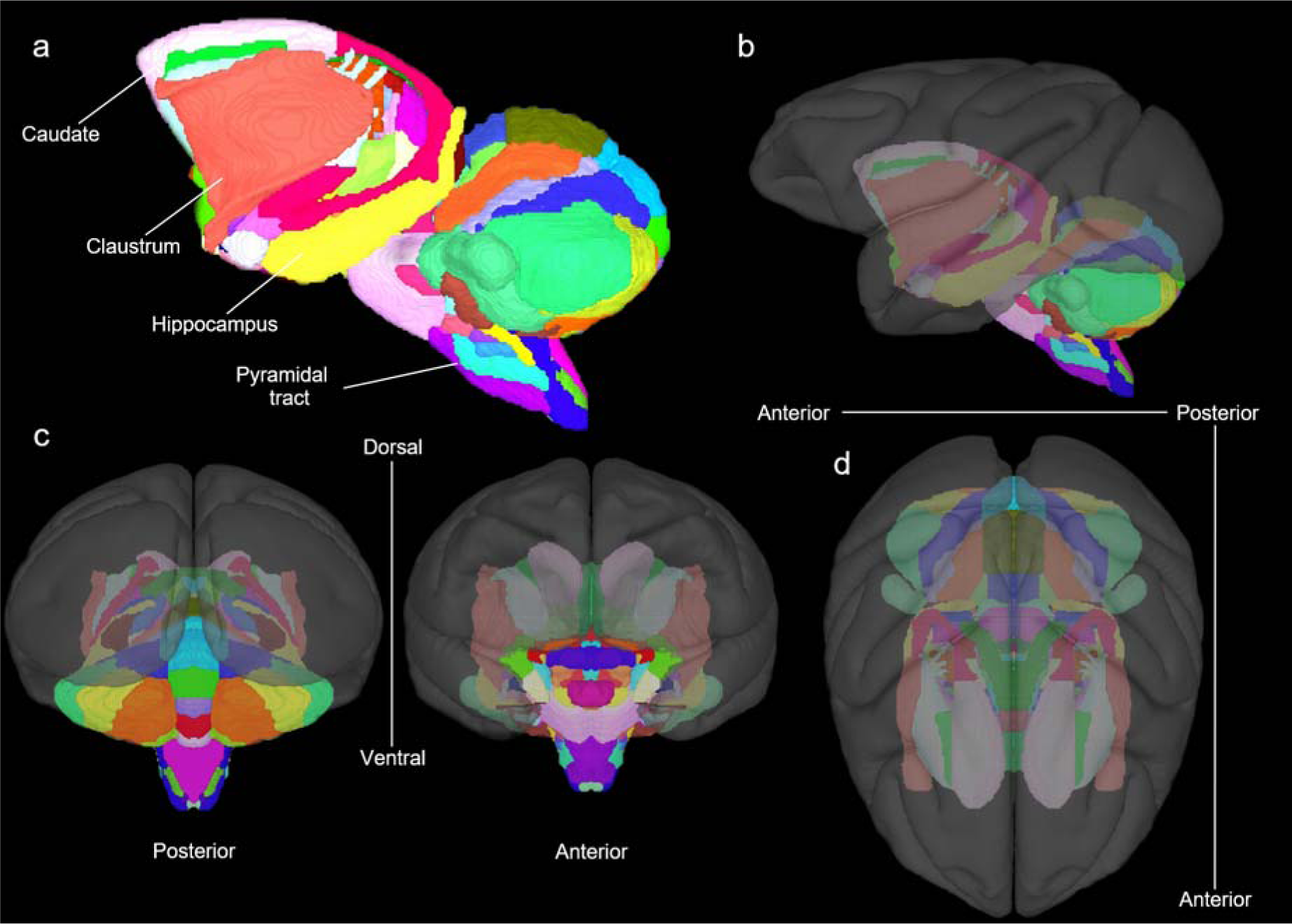
Surface views of SARM (level 6) in the NMT v2.0 symmetric template. Volumetric atlas regions were converted into individual surfaces for surface-based analysis. (**a**) A lateral view of the subcortical surfaces, displayed in color using SUMA (Saad et al., 2004). The subcortical regions are shown with respect to the NMT v2 surface (shown in gray scale) in a (**b**) left lateral, (**c**) posterior (left), anterior (right), and (**d**) superior view.

**Figure 3.**
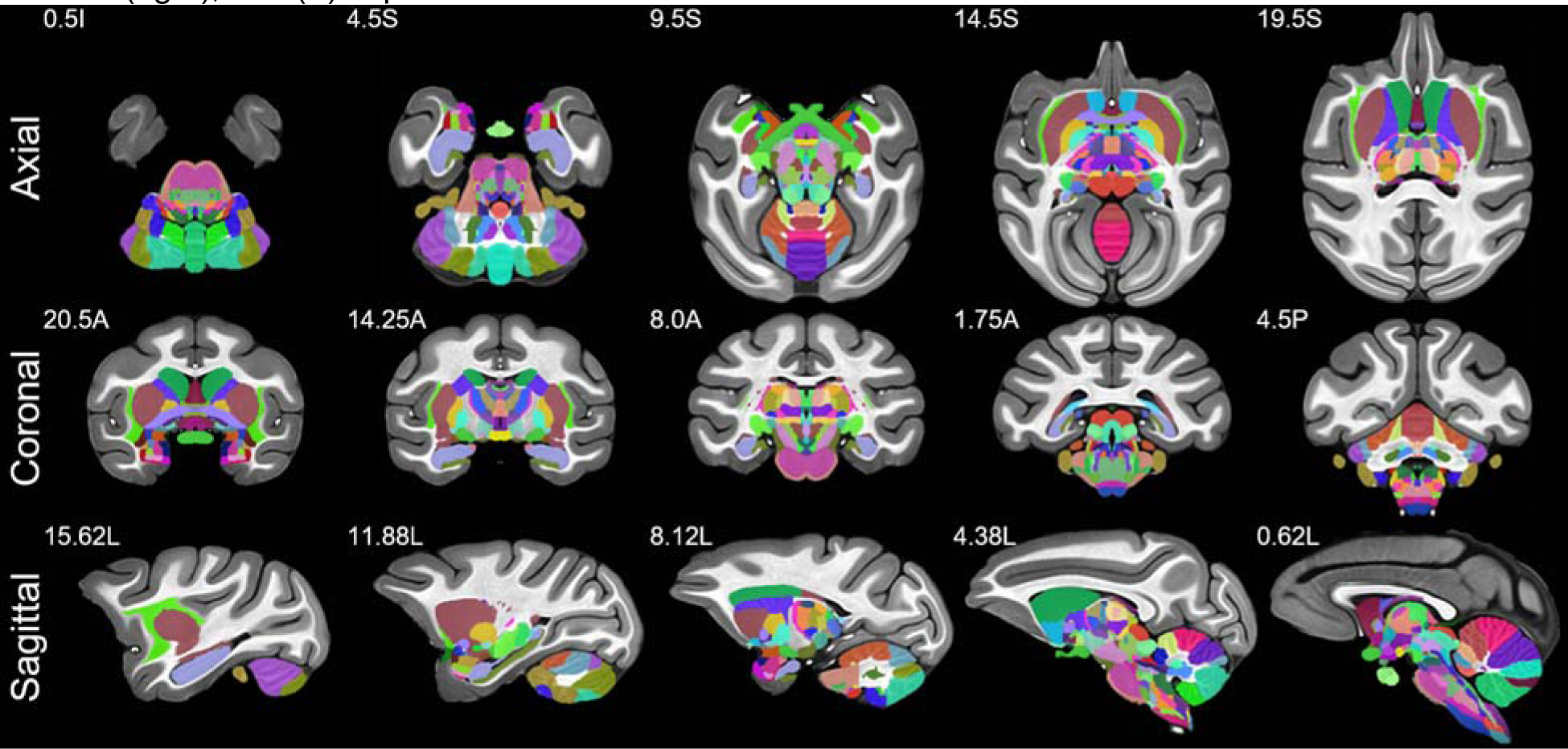
Subcortical regions in NMT v2 space. The subcortical parcellation of the G12 subject was warped to the NMT v2 symmetric template and manually adjusted to match the template’s morphology. These regions constitute level 6 of the SARM and are shown in color on the symmetric brain-extracted NMT v2 template. Slice coordinates relative to the origin (EBZ; ear bar zero) are in mm in the superior/inferior (top), anterior/posterior (middle), and left/right (bottom) directions.

#### 3.1.2. Hierarchical Grouping

The 206 manually-drawn primary ROIs comprise the finest level (level 6) of the SARM. Following the same principle as in the CHARM (Jung et al., this issue), these 206 ROIs were organized hierarchically into six levels of granularity. Individual ROIs were assembled into progressively larger (and, in most cases, spatially contiguous) groups from levels 5 to 1. Each ROI or group of ROIs at a lower level (e.g., level 4) belongs to exactly one group in the next higher level (e.g., level 3). Table 1 shows the number of ROIs in each level and characterizes their volumes in the NMT v2. Whole-brain coronal views of the SARM levels 2, 4, and 6 are shown in Figure 4. The various levels were designed to be suitable for either structural or functional MRI analyses, with their different spatial resolutions. Users can combine more than one grouping level within a single analysis to, for instance, examine the relationships between a specific nucleus and larger composite brain regions. To further illustrate the SARM hierarchy, Figure 5 provides an exploration of the amygdala. The dendrogram demonstrates how the amygdala splits into its constituent regions, and these component structures are depicted on a coronal section for levels 3-6.

**Figure 4.**
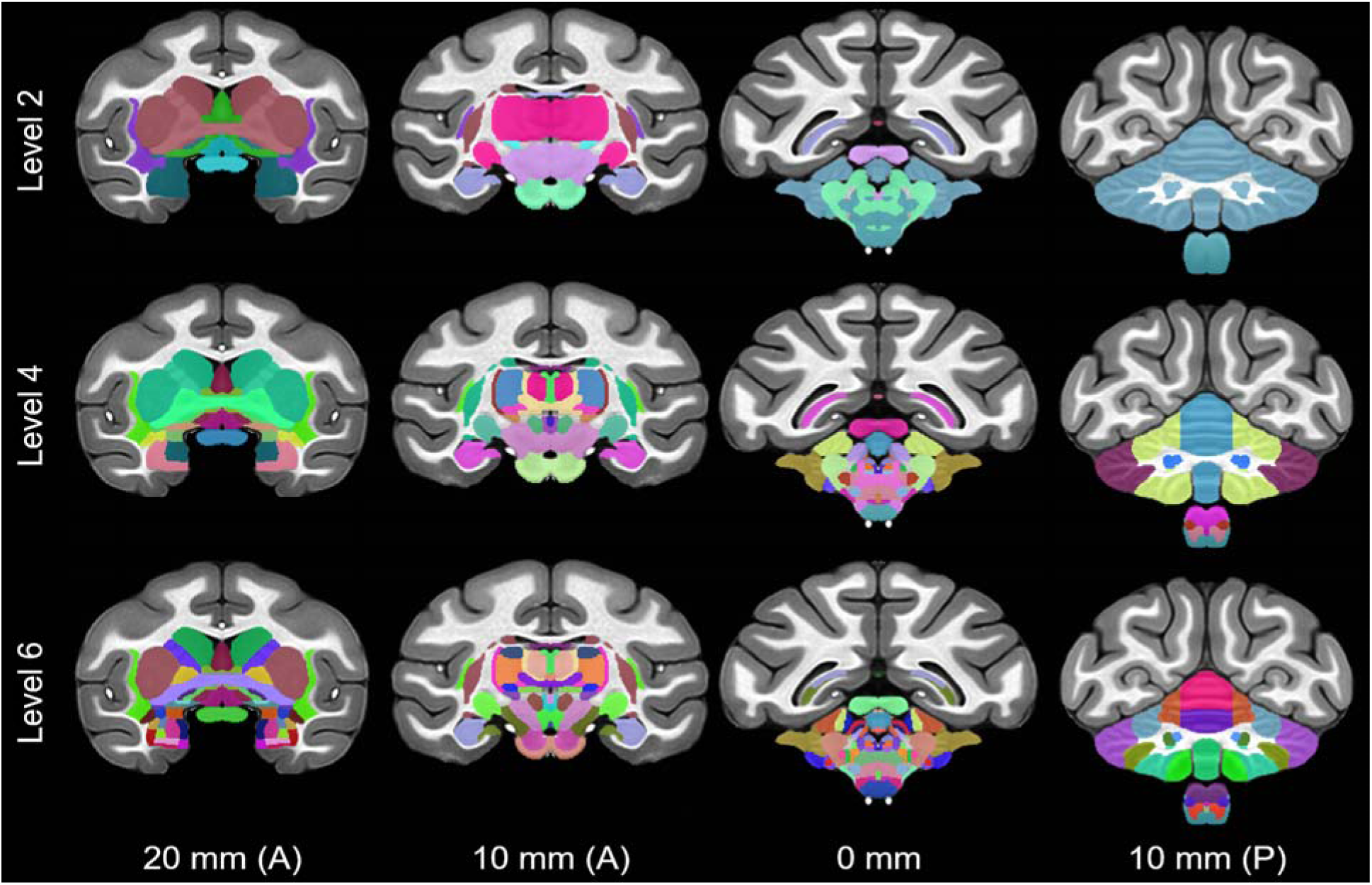
SARM’s Hierarchical ROI Groupings. Representative coronal slices through the symmetri NMT v2 in stereotaxic space, showing levels 2, 4 and 6 of the SARM hierarchy. Level 2 contains relatively broad composite structures, level 4 contains somewhat finer groupings, and level 6 consists of the finest anatomical segmentation. Slice coordinates are in mm anterior (A) or posterior (P) to the origin (EBZ; ear bar zero).

**Figure 5.**
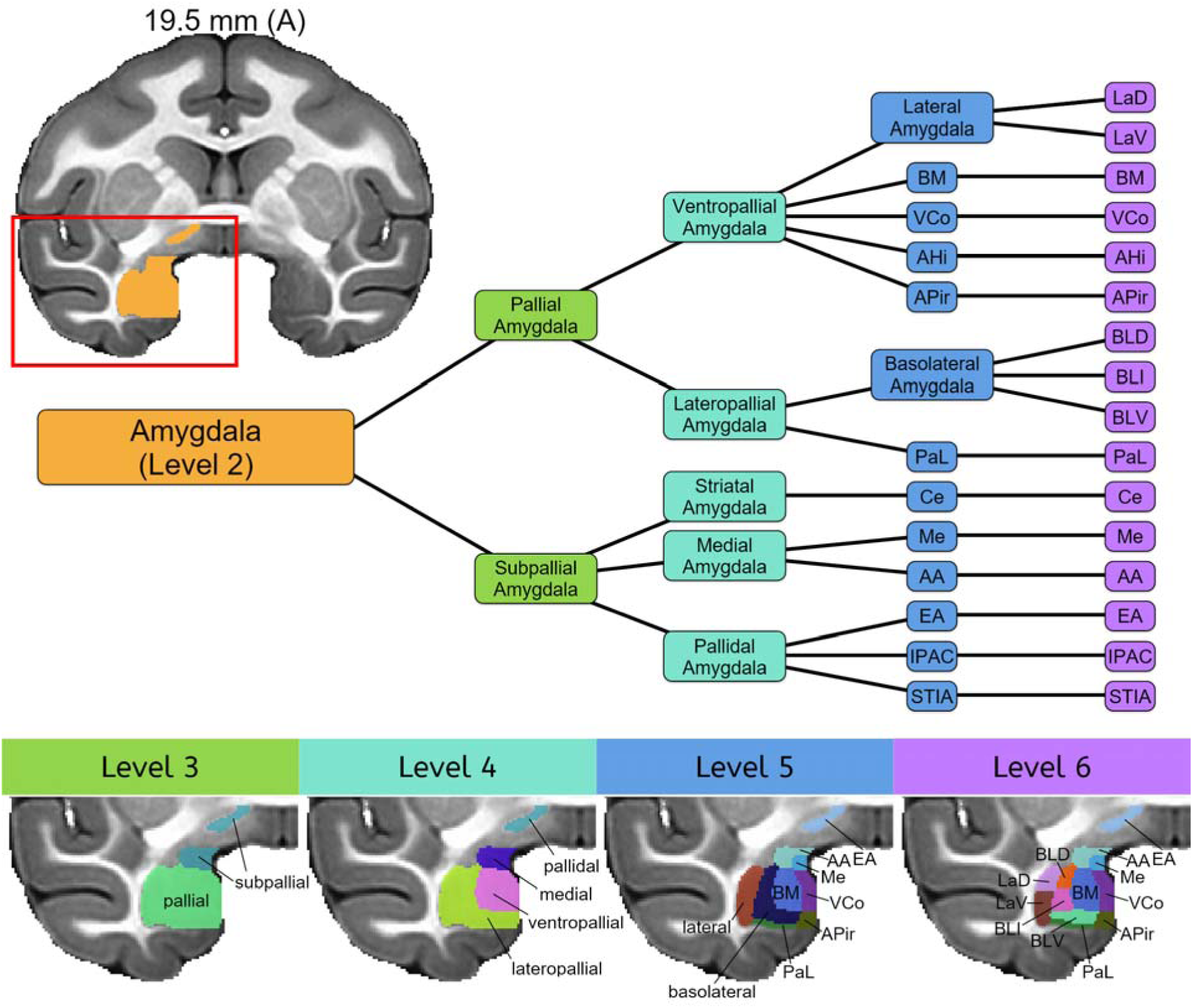
Hierarchical parcellation of the amygdala. The amygdala, a subcortical region within the telencephalon, is shown in orange on the right hemisphere of a coronal section. The bottom row shows amygdala subdivisions for levels 3-6 of the hierarchical atlas in various colors on close ups of the right temporal lobe region contained in the red box on the full coronal section. The color-coded dendrogram shows the hierarchical relationship between the components of levels 3-6. Level 3 distinguishes the portions of amygdala deriving from the pallium and subpallium during development. At level 4, regions arising from particular domains within the pallium or subpallium are differentiated. These regions are further divided into the various amygdala nuclei and subnuclei in levels 5 and 6. Note other subcortical regions are not shown. ***Abbreviations***: ***AA***, anterior amygdaloid area; ***AHi***, amygdalohippocampal area; ***APir***, amygdalopiriform transition; ***BLD***, ***BLI***, and ***BLV***, basolateral dorsal, intermediate, and ventral amygdaloid n.; ***BM***, basomedial amygdaloid n.; ***Ce***, central amygdaloid n.; ***EA***, extended amygdala; ***IPAC***, posterior interstitial nucleus; ***LaD*** and ***LaV***, lateral dorsal and ventral amygdaloid n.; ***Me***, median amygdaloid n.; ***PaL***, paralaminar amygdaloid n.; ***STIA***, intraamygdaloid division of the bed n. of the stria terminalis; ***VCo***, ventral cortical amygdaloid n..

**Table 1.**
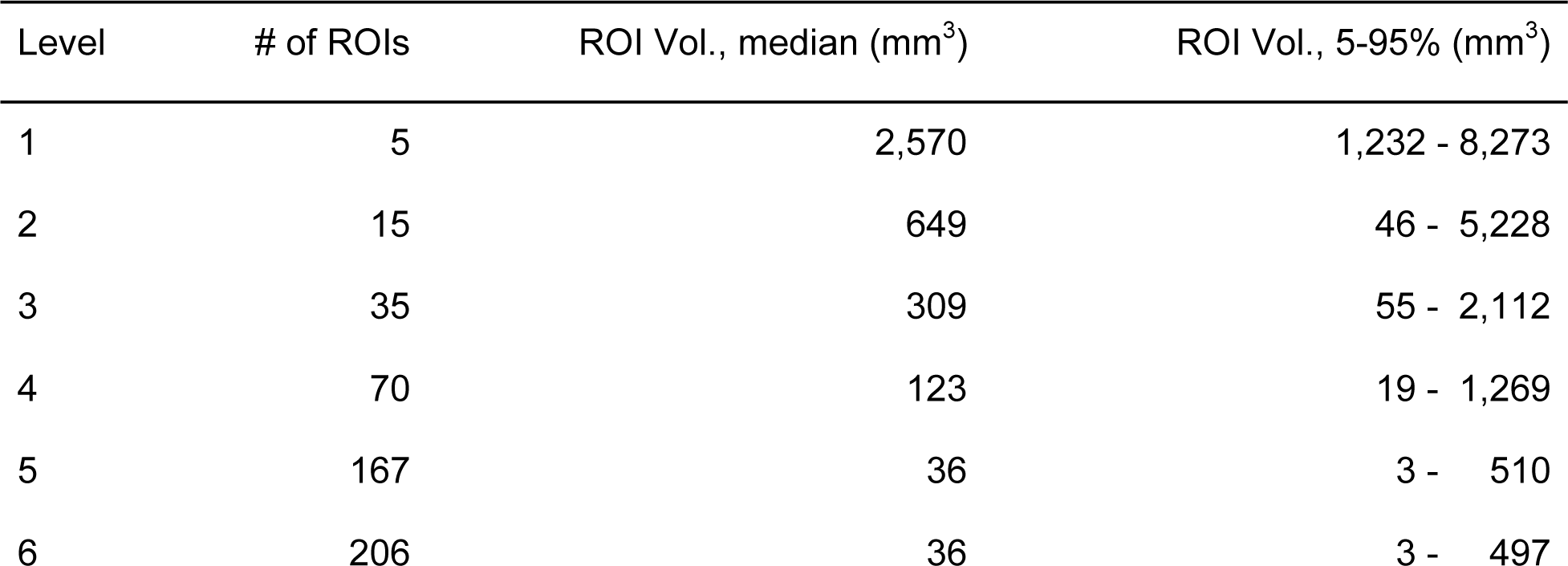
The SARM Group Hierarchy. For each level of the hierarchy, the number of ROIs used to parcellate the subcortex, their median volume, and the 5^th^-95^th^ percentile of their volumes are listed. At lower levels, ROIs are combined into fewer and larger composite structures. The full table of SARM region names, abbreviations, and constituents is provided as a CSV file in the distribution package and i also shown in Table S1.

| Level | # of ROIs | ROI Vol., median (mm<sup>3</sup>) | ROI Vol., 5-95% (mm<sup>3</sup>) |
| --- | --- | --- | --- |
| 1 | 5 | 2,570 | 1,232 - 8,273 |
| 2 | 15 | 649 | 46 - 5,228 |
| 3 | 35 | 309 | 55 - 2,112 |
| 4 | 70 | 123 | 19 - 1,269 |
| 5 | 167 | 36 | 3 - 510 |
| 6 | 206 | 36 | 3 - 497 |

#### 3.1.3 ROI and Hierarchical Grouping Definition

Supplementary Table 1 (Table S1) itemizes the 206 ROIs at level 6 and shows their progressive hierarchical grouping, from level 5 to level 1. Level 1 assembles all ROIs according to the classical developmental division of the neuraxis, namely the (subcortical) telencephalon, diencephalon, mesencephalon, metencephalon, and myelencephalon. The ordering of these ROIs reflects that, together, the subcortical telencephalon and diencephalon comprise the subcortical forebrain, the mesencephalon is synonymous with the midbrain, and the metencephalon and myelencephalon make up the hindbrain. Level 2 divides the telencephalon into lateral and ventral pallium (LVPal), medial pallium (MPal), amygdala (Amy), basal ganglia (BG), diagonal subpallium (DSP), and preoptic (preoptic) regions. The order in which these divisions are listed roughly follows the developmental partition proposed by Puelles et al. (2013), with entirely pallial groups (LVPal and MPal) first, followed by the amygdala with its pallial and subpallial components (see level 3), and, finally, by entirely subpallial groups (BG, DSP and preoptic). At level 2, the diencephalon is divided into the hypothalamus (Hy), prethalamus (PreThal), thalamus (Thal), and epithalamus (EpiThal). The mesencephalon was not divided at level 2, but the region was relabeled “midbrain” (Mid) to match the more common choice of terminology employed beyond level 2. Still at level 2, the metencephalon was split into the pons (Pons) and the cerebellum (Cb), whereas the myelencephalon remained whole, but switched names to the term medulla (Med). Levels 3 to 6 propose a progressively more refined parcellation of the larger groups of level 2, ending with level 6, which lists each individually drawn ROI. Levels 5 and 6 were left largely similar to allow for future versions of the SARM (now SARM v1) to incorporate additional sub-structures. In general, beyond level 2, the hypothalamic, thalamic, mes-, met- and myel-encephalic ROIs were not grouped according to the ontological plan because most of the small ontologically related ROIs of these regions are spatially non-contiguous in the adult brain. Instead, these ROIs were mainly grouped according to either functional or purely topological criteria, with the practical condition that they remain contiguous, as this has greater relevance for targeting of subcortical regions and neuroimaging analytical strategies (e.g. clustering). The next sections briefly describe the rationale for the drawing of the ROIs and their grouping at and below level 2.

##### 3.1.3.1 Lateral and Ventral Pallium

The lateral and ventral pallium (LVPal) group contains 4 primary ROIs (Table S1). The claustrum (Cl) and the dorsal and ventral endopiriform claustrum (DEn and VEn) were all identifiable in the G12 (not shown) and the NMT v2 (Fig. 6a,b). DEn appeared as a separate entity at the ‘heel’ of Cl (see arrows in Fig. 6a). VEn was recognized by a consistently lighter contrast in comparison with the darker nuclei of the amygdala (see blue asterisk in Fig. 6a). The small bundle of capillaries (base of the lenticulostriatal arteries) located at the base of the putamen (Pu) and above the ‘heel’ of Cl was incorporated into the Pu ROI by default (yellow asterisk in Fig. 6a). The piriform cortex (Pir; not shown) was recognized by its thinner cortical width at the medial junction of the orbitofrontal and temporal cortices (Carmichael and Price, 1994; Evrard et al., 2014), although its exact border with Cl at the limen insulae and with the amygdalo-piriform transition (APir) in the temporal lobe could not be ascertained.

**Figure 6.**
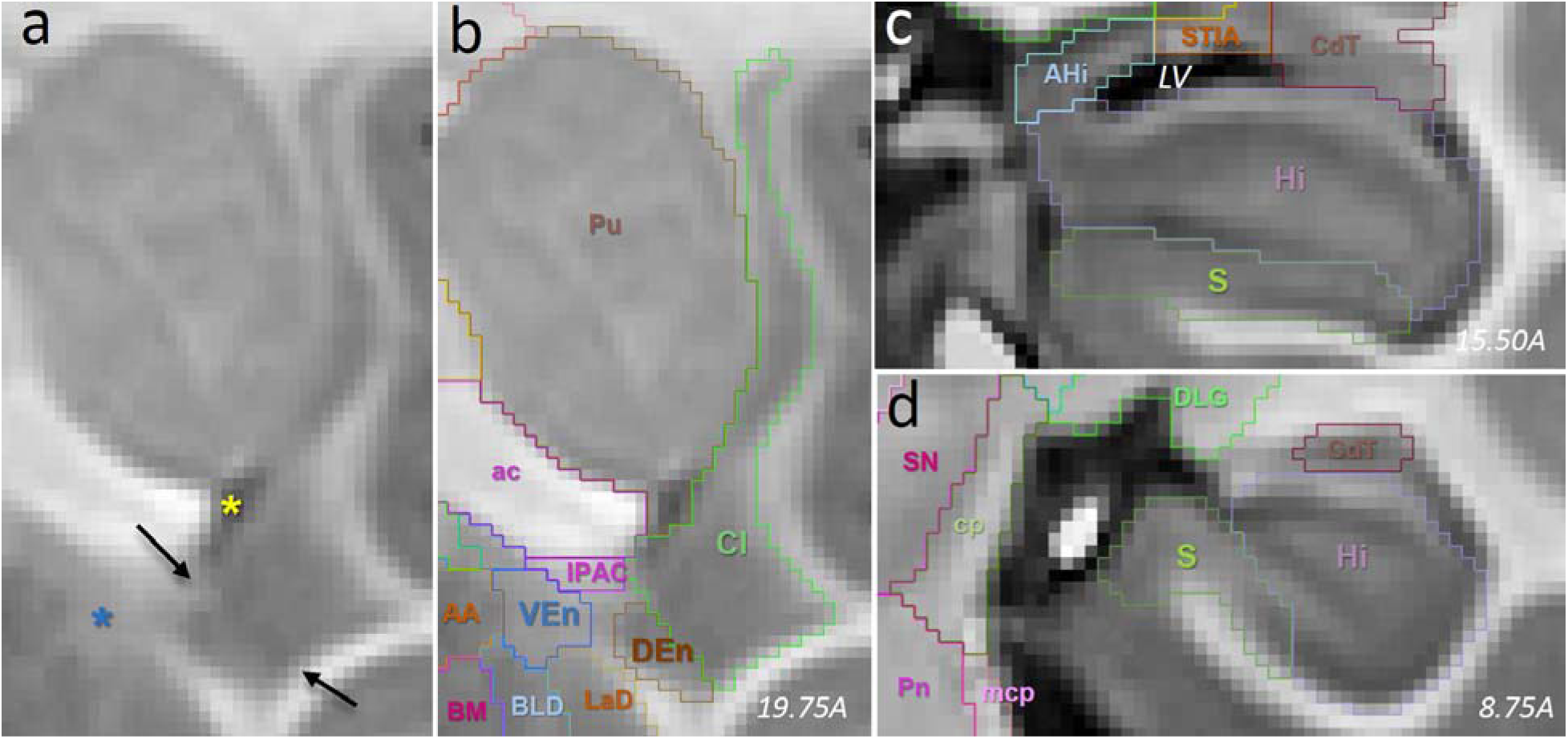
SARM lateral, ventral, and medial pallial ROIs. (**a,b**) Coronal section through the right hemisphere of NMT v2 (corresp. to RMBSC Fig. 48) with delineations of Cl, DEn and VEn. In (**a**), the black arrows indicate the border between Cl and DEn; the blue asterisk marks the consistently lighter contrast of VEn; the yellow asterisk indicates a bundle of capillaries incorporated into the Pu ROI. (**c**,**d**) Coronal sections through the right hemisphere of NMT v2 (RMBSC4 Fig. 63 and 77, respectively) illustrating the delineations of Hi and S. ***Abbreviations***: ***AA***, anterior amygdaloid area; ***ac***, anterior commissure; ***AHi***, amygdalohippocampal area; ***CdT***, tail of the caudate; ***BM***, basomedial amygdaloid n.; ***BLD***, basolateral amygdaloid n.; ***cp***, cerebral peduncle; ***Cl***, claustrum; ***DEn***, dorsal endopiriform claustrum; ***DLG***, dorsal lateral geniculate; ***Hi***, hippocampus; ***IPAC***, interstitial nucleus of the posterior part of the anterior commissure; ***LV***, lateral ventricle; ***mcp***, medial cerebellar peduncle; ***Pn***, pontine nucleus; ***Pu***, putamen; ***S***, subicular complex; ***SN***, substantia nigra; ***STIA***, intraamygdaloid stria terminalis; ***VEn***, ventral endopiriform claustrum. In all panels, left is medial and top is dorsal.

At level 4, DEn and VEn are grouped in En (Table S1). At level 3, En is grouped with Pir in the ventral pallium (VPal), from which En and Pir originate along with other olfactory structures (Puelles et al., 2013) that were not segmented here (but see CHARM; Jung et al., this issue). Also at level 3, Cl constitutes the only ROI of the lateral pallium (LPal). At level 2, LPal and VPal are grouped under LVPal.

##### 3.1.3.2 Medial Pallium (Hippocampal Formation)

The hippocampus (Hi) and subicular complex (S) were segmented without distinguishing their internal subdivisions (i.e., CA1-3, dentate gyrus, parasubiculum, presubiculum, subiculum *per se*, and prosubiculum) (Table S1; Fig. 6c,d). They were grouped together, along with the fimbria (fi), as hippocampal formation (HF) at levels 3 and 4. Although being acellular, the fimbria was added to the HF group in order to take into account the lower spatial resolution of functional scans that may not distinguish fi from Hi and S. The fornix (f) was drawn throughout the lateral ventricle mainly for illustrative purposes. It was not added to the HF group to avoid false attribution of activation possibly originating from regions located in the vicinity of the distant f (e.g., septum and dorsal thalamus). Finally, at level 2, HF and f were grouped together in the medial pallium (MPal), from which they originate (Puelles et al., 2013).

##### 3.1.3.2 Amygdala

The amygdala was delineated into 16 primary ROIs (Table S1). Figure 7 illustrates the segmentation of the amygdala at one representative anteroposterior level in NMT v2, G12 and RMBSC4. Throughout the anteroposterior extent of the amygdala, the dorsal and ventral parts of the lateral amygdaloid nucleus (LaD and LaV) were recognizable by their darker contrast, compared to the lighter dorsal, intermediate and ventral parts of the basolateral nucleus (BLD, BLI and BLV). The theoretical location of the basomedial nucleus (BM) often contained a darker region in both NMT v2 and G12, which likely corresponds to the parvocellular or magnocellular division of BM and contrasts with the lighter and more homogeneous ventral cortical nucleus (VCo). The paralaminar nucleus (PaL) appeared as a thin sheet of lighter (G12), and somewhat darker (NMT v2), contrast at the base of the amygdala. The central nucleus (Ce) was less distinct, but its theoretical anatomical location largely corresponded to a circular area with a lighter contrast in NMT v2. Medial to Ce, AA and the medial nucleus (Me) appeared darker in NMT v2 and lighter in G12. The boundaries between Me and AA, between BLD, BLI, and BLV, and between LaD and LaV were drawn based on their theoretical topological localization (Amaral et al., 1992; Stefanacci et al., 2000; Paxinos et al., 2009). Lateral to the amygdala, the amygdalostriatal transition area (ASt) appears as a distinct region, separated by a thin but distinct lighter (NMT v2) or darker (G12) strip of white matter. Dorsal to the amygdala proper, the interstitial nucleus of the posterior part of the anterior commissure (IPAC), the intra-amygdaloid division of the bed nucleus of the stria terminalis (STIA), and the extended amygdala (EA) are all readily distinguishable in NMT v2. For example, EA appears as a distinct lighter band underneath the ventral pallidum (VP) (Fig. 8a).

**Figure 7.**
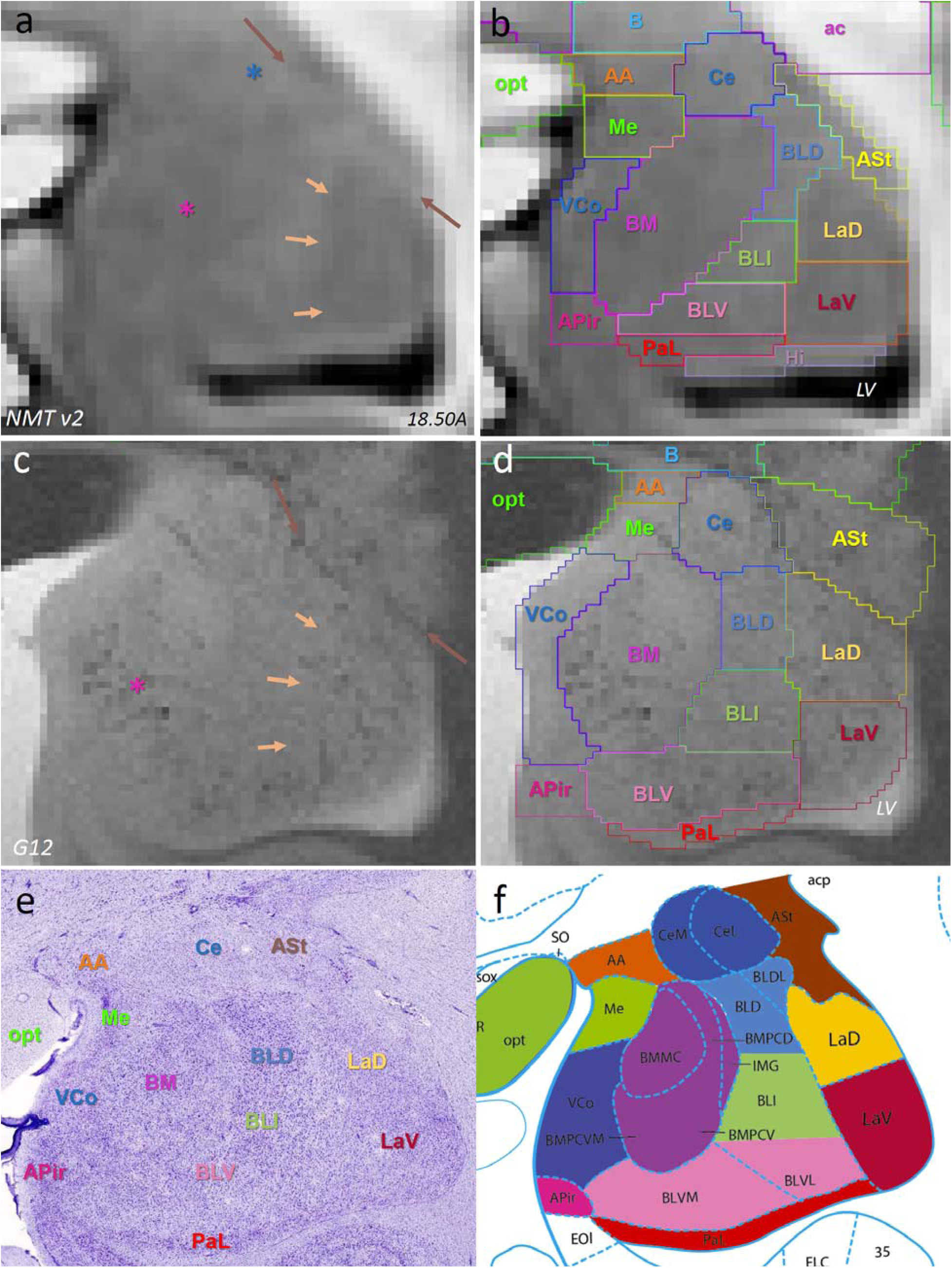
SARM’s amygdaloid ROIs. Coronal slices through the right hemisphere at approximately the same location in (**a**, **b**) NMT v2, (**c**, **d**) G12, and (**e**,**f**) a corresponding pair of RMBSC4 Nissl-stained section and diagram (Fig. 56). The brown arrow in (**a**,**c**) points at the boundary between ASt and the amygdala. The blue asterisk in (**a**) points at a zone of lighter contrast, likely corresponding to Ce. The light orange arrows in (**a**, **c**) point at the putative border of LaD and LaV with BLD, BLI and BLV. The mauve asterisks in panels (**a**, **c**) mark the darker contrast included in BM. ***Abbreviations***: ***AA***, anterior amygdaloid area; ***ac***, anterior commissure; ***APir***, amygdalopiriform transition area; ***ASt***, amygdalostriatal transition area; ***B***, basal n.; ***BLD***, ***BLI***, ***BLV***, dorsal, intermediate and ventral parts of the basolateral amygdaloid n.; ***BM***, basomedial amygdaloid n.; ***Ce***, central amygdaloid n.; ***LaD*** and ***LaV***, dorsal and ventral parts of the lateral amygdaloid n.; ***LV***, lateral ventricle; ***Me***, medial amygdaloid n.; ***opt***, optic tract/chiasma; ***PaL***, paralaminar amygdaloid n.; ***VCo***, ventral cortical amygdaloid n.. In all panels, left is medial and top is dorsal.

**Figure 8.**
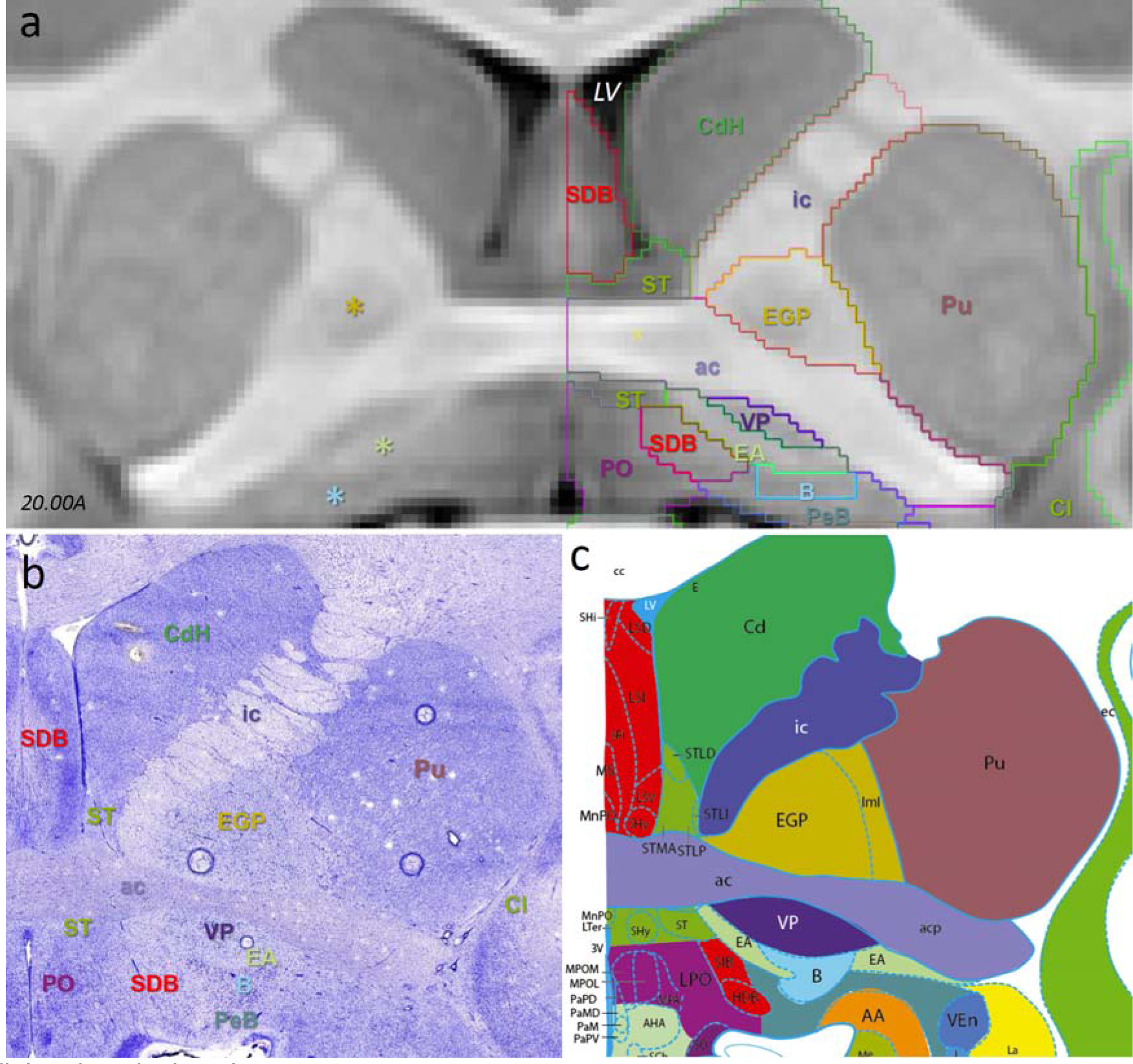
SARM telencephalic ROIs. Coronal slices through (**a**) the symmetric NMT v2 and (**b**,**c**) the approximately corresponding pair of RMBSC4 Nissl-stained section and diagram (Fig. 50). The asterisks on the left side in (**a**) emphasize contrasts corresponding to the location of EGP (yellow), EA (pale green), and B (blue). Only the anterior portion of SFi is included in the SDB ROI; further caudally, SFi is recognized as a single ROI (see Fig. 9). In all panels, the top is dorsal. In (**b**,**c**), left is medial. ***Abbreviations***: ***B***, basal nucleus; ***CdH***, head of the caudate nucleus; ***Cl***, claustrum; ***EA***, extended amygdala; ***EGP***, external globus pallidus; ***ic***, internal capsule; ***PeB***, peri-basal region; ***PO***, preoptic area; ***Pu***, putamen; ***SDB***, septum and diagonal band; ***ST***, stria terminalis; ***VP***, ventral pallidum. For the missing abbreviations in (**c**), see Paxinos et al., 2009, where most abbreviations are similar to Paxinos et al., in preparation.

As shown in Fig. 5, the amygdala is divided at level 3 on developmental grounds into its pallial and subpallial portions. At level 4, the former splits into the portions arising from the ventral and lateral pallium, while the latter divides into the striatal, medial, and pallidal amygdala. These are then further divided into individual nuclei (level 5) and, in the case of the lateral and basolateral nuclei, into subnuclei (level 6). The different ROIs of the pallidal amygdala are not all contiguous. For example, EA has no boundary with other amygdaloid nuclei. Therefore, the pallidal amygdala ROI is one of the two SARM group ROIs containing non-contiguous ROIs. See 3.1.3.11 for the other exception in the medulla.

##### 3.1.3.4 Basal Ganglia

The basal ganglia (BG) was delineated into 12 primary ROIs (Table S1). The head of the caudate (CdH) and the putamen (Pu) were identifiable due to their prominent size and distinct boundary with the internal capsule (ic), anterior capsule (ac), corpus callosum (cc), external capsule (ec), and white matter of the cerebral cortex (Fig. 8). The ventral boundary between CdH and the anterior portion of the bed nucleus of the stria terminalis (ST) was marked by an abrupt darkening of the signal in ST (see also Section 3.1.3.5). The ventral boundaries of CdH and Pu with the accumbens nucleus (Acb; not shown) were identified by a consistent change to a more heterogeneous contrast pattern in Acb. The tail of the caudate (CdT) was distinct all along the lateral ventricle (LV) (Fig. 10) and in proximity to the amygdala, where it borders the amygdalostriatal transition area (ASt; not shown). The external (EGP; Fig. 8) and internal (IGP; not shown) globus pallidus were readily identifiable due to their slightly darker contrast, compared to the surrounding white matter. They were distinguishable from one another due to their characteristic shapes and their separation by the thin medial medullary lamina (not shown). The ventral pallidum (VP) appeared distinctly darker between ac and EA (Fig. 8).

**Figure 10.**
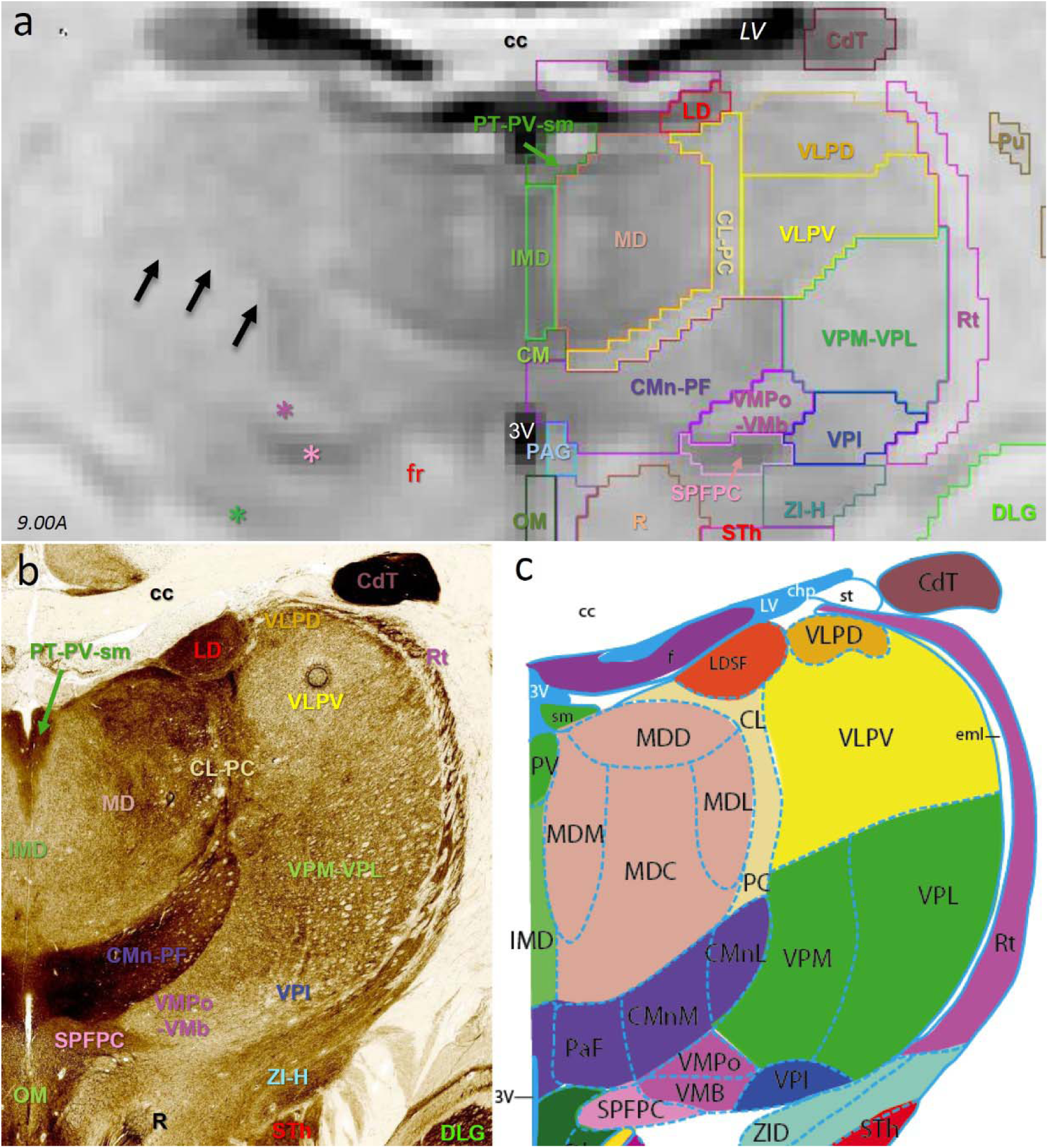
SARM’s thalamic ROIs in coronal view. (**a**-**c**) Coronal slice through the NMT v2 in stereotaxic space (**a**), and a corresponding pair of RMBSC4 Nissl AChE stain section (**b**) and diagram (**c**) (Fig. 71). In (**a)**, the black arrows point at the boundary between VLPV and VPM-VPL. The asterisks emphasize the localizations of VMPo-VMb (dark pink), SPFPC (light pink), and ZI-H (green). The red “fr” indicates the localization of the fasciculus retroflexus, ventral to CMn-PF. (In more posterior slices, fr ascends through CMn-Pf, as illustrated in Fig. 9b and 11a.) ***Abbreviations*** *(**a**,**b**)*: ***3v***, third ventricle; ***CdT***, the tail of the caudate; ***CL-PC***, centrolateral and paracentral thal. n.; ***CM***, central medial thal. n.; ***CMn-PF***, centromedial and parafascicular thal. n.; ***DLG***, dorsolateral geniculate thal. n.; ***IMD***, intermediodorsal thal. n.; ***LD***, laterodorsal thal. n.; ***LV***, lateral ventricle; ***MD***, mediodorsal thal. n.; ***OM***, oculomotor complex; ***PAG***, periaqueductal gray; ***R***, red n.; ***Rt***, reticular thal. n.; ***Pu***, putamen; ***PT-PV-sm***, ensemble of the stria medullaris, paraventral nucleus and paratenial thal. n.; ***SPFPC***, subparafascicular parvocellular thal. n.; ***STh***, subthalamic n.; ***VLPD*** and ***VLPV***, posterodorsal and posteroventral parts of the ventrolateral thal. n.; ***VMPo-VMB***, posterior and basal parts of the ventromedial thal. n.; ***VPI***, ventroposterior inferior thal. n.; ***VPM-VPL***, ventroposterior medial and lateral thal. n.; ***ZI-H***, zona incerta and lenticular fascicles (H fields). For the missing abbreviations in (**c**), see Paxinos et al., 2009, where most abbreviations are similar to Paxinos et al., in preparation.

At level 5, CdH and CdT were grouped as caudate (Cd), and EGP, IGP and their ventral ansa lenticularis tract (al; not shown) were grouped as globus pallidus (GP). At level 4, Cd, Pu, ASt, and ic (which contains strands of neurons) were grouped as dorsal striatum (DStr), and Acb and Tu were grouped as ventral striatum (VStr). At level 3, DStr and VStr were grouped as striatum. GP, ac and VP were grouped as pallidum (Pd) at levels 4 and 3. The inclusion of white matter tracts (e.g., ac in Pd) enables using broad ROIs in fMRI analyses with rather low spatial resolution, and in which the BOLD signal would likely ‘spread’ over multiple neighboring structures, without possible distinction between smaller ROIs.

##### 3.1.3.5 Diagonal Subpallium

The ontological definition of the diagonal subpallium (DSP) includes the basal nucleus of Meynert (B), the bed nucleus of the stria terminalis (ST), and different parts of the septum and diagonal band of Broca region (SDB and SFi) (Table S1) (Puelles et al., 2013). B is anatomically formed by an ill-defined group of cholinergic neurons at the base of the basal ganglia. In some slices of NMT v2, the putative location of B could correspond to a slightly darker region ventral to EA and VP (see blue asterisk in Fig. 8a, left); however, this appearance is not consistent. Thus, for the most part, the delineation of B is based on its most likely localization, underneath VP anteriorly (Fig. 8) and in between IGP, the optic tract (opt) and Pu posteriorly (not shown; see for example RMBSC4 Fig. 66). To take into account this less obvious delineation, the region surrounding our delineation of B is labeled as ‘peri-basal region’ in the SARM v1 (PeB; Fig. 8), which corresponds, in RMBSC4 to a rather undefined region sandwiched between B and other regions such as AA, Ce and SDB. B and PeB are grouped in the basal nucleus ‘region’ (BR) at levels 5 and 4. Anteriorly, the different parts of ST form a distinct ‘ring’ of dark signal around ac (ST; Fig. 9a,b). Posteriorly, ST mingles with various fiber tracts and appears lighter (Fig. 9a,b). The different components of the medial and lateral septum, as well as those of the diagonal band of Broca, were not readily distinguishable from one another in G12, although the medial portion of the septum appears lighter in NMT v2 and could be ascribed to the medial septum in a future version of the SARM (Fig. 8). Ventrally and posteriorly, SDB (i.e., SIB and HDB in Figure 8c) is consistently darker than EA but lighter than PO, and sits ‘sandwiched’ between them. For the time being, these anterior and posteroventral regions are grouped in SDB. However, dorsally and posteriorly, one component of the septum that runs along the anterior portion of the lateral ventricle, namely the septimbrial nucleus (SFi), was labeled as a distinct ROI (Fig. 9). SDB and SFi are grouped in the septum diagonal band region (SDBR) at level 4. BR, ST, and SDBR are grouped in DSP at levels 3 and 2.

**Figure 9.**
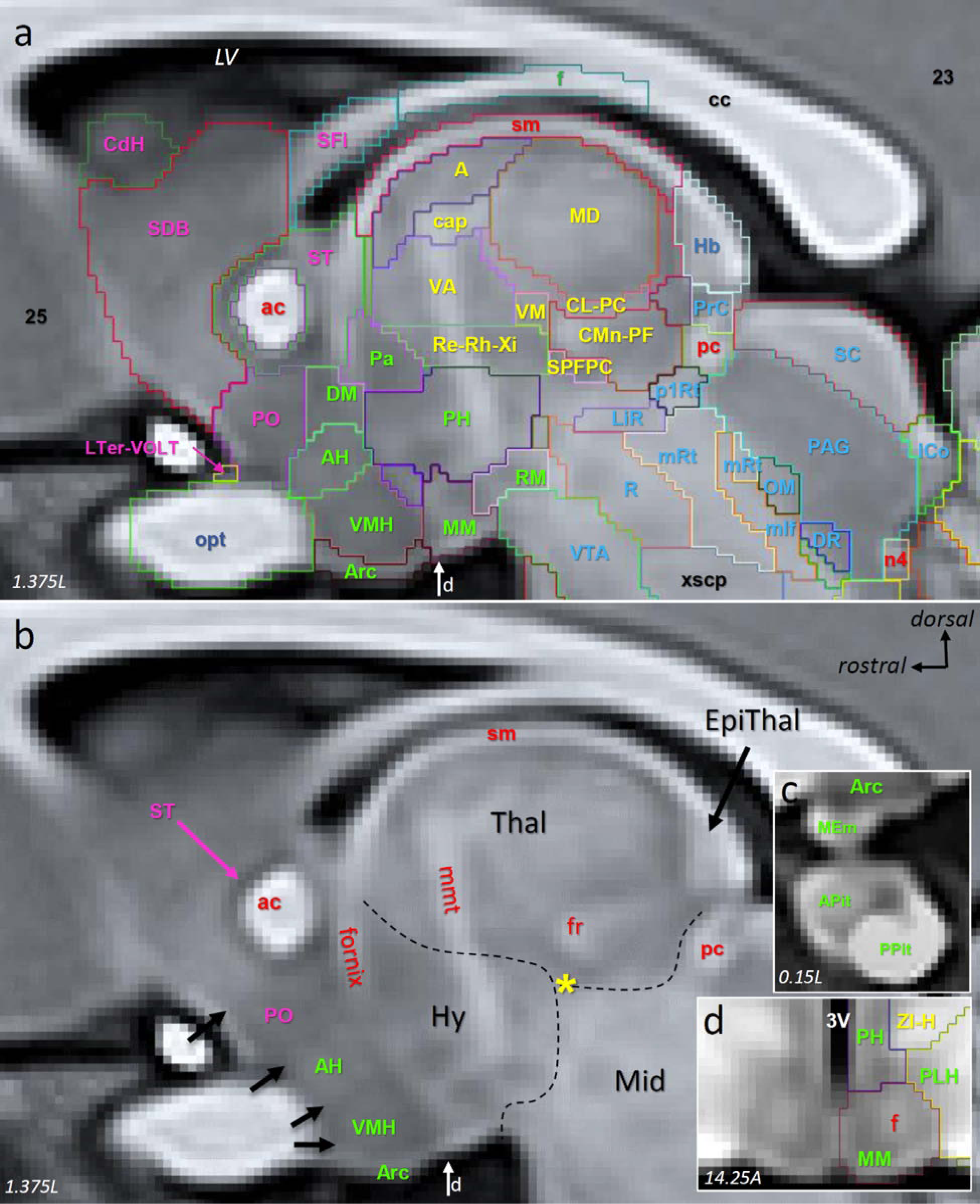
SARM ROIs in parasagittal view. (**a, b**) Sagittal slice through the NMT v2 showing (**a**) the delineations of SARM regions and (**b**) major anatomical landmarks. In (**a**), ROI labels are color coded: telencephalic (magenta), hypothalamic (green), thalamic (yellow), epithalamic (dark blue), midbrain (light blue), and pons (black). Notable landmarks in (**b**) include the fr, mmt (not a ROI, inserted in figure for orientation purpose) and fornix. The yellow asterisk is placed just below the distinct darker contrast that characterizes SPFPC (see also Fig. 10). The thin dashed lines emphasize the distinctive change in contrast between Thal, Mid, and Hy. The pink arrow points at the ring of dark contrast of ST around ac. The four black arrows on the left side of (**b**) mark the contrast changes between PO, AH, VMH, and Arc. Panel (**c**) shows a mid-sagittal view of the subjacent (ventral to Arc) pituitary regions APit and PPit, as well as MEm. Panel (**d**) shows a symmetrical coronal view of MM with a distinctively lighter contrast in its center, corresponding to f. The vertical white arrow at the base of the hypothalamus in panels (**a**,**b**) indicates the anteroposterior level of the coronal view shown in panel (**d**). ***Abbreviations***: ***23*** and ***25***, cortical areas 23 and 25; ***3V***, third ventricle; ***A***, anterior thal. n.; ***ac***, anterior commissure; ***AH***, anterior hy. n.; ***APit***, anterior pituitary; ***Arc***., arcuate n.; ***cap***, capsule of the anterior thalamic nucleus; ***cc***, corpus callosum; ***CdH***, head of the caudate nucleus; ***CL-PC***, centrolateral and paracentral thal. n.; CMn-PF, centromedian and parafascicular thal. n.; ***DM***, dorsomedial hy. n.; ***DR***, dorsal Raphe; ***EpiThal***, epithalamus; ***f***, fornix; ***fr***, fasciculus retroflexus; ***Hb***, habenula; ***Hy***, hypothalamus; ***ICo***, inferior colliculus; ***LH***, lateral hypothalamic area; ***LiR***, linear Raphe; ***LTer-VOLT***, lamina terminalis and vascular organ of the lamina terminalis; ***LV***, lateral ventricle; ***MD***, mediodorsal thal. n.; ***MEm***, medial eminence, ***Mid***, midbrain; ***mlf***, medial longitudinal fascicle; ***MM***, mammillary n.; ***mRt*,** midbrain reticulum; ***n4***, 4^th^ cranial nerve (crossing); ***OM***, oculomotor complex; ***p1RT*,** prosomere 1 reticulum; ***Pa***, paraventricular hy. n.; ***PAG***, periaqueductal gray; ***pc***, posterior commissure; ***PH***, posterior hy. n.; ***PLH***, peduncular lateral hypothalamus; ***PO***, preoptic area; ***PPit***, posterior pituitary; ***PrC***, precommissural n.; ***R***, red n.; ***Re-Rh-Xi***, reuniens, rhomboid and xiphoid thal. n.; ***RM***, retro-mammillary n.; ***SC***, superior colliculus; ***SDB***, septum-diagonal band; ***SFi***, septimbrial n.; ***sm***, stria medullaris; ***SPFPC***, subparafascicular parvocellular thal. n.; ***ST***, bed n. of the stria terminalis; ***Thal***, thalamus; ***VA***, ventral anterior thal. n.; ***VM***, ventromedial thal. n.; ***VMH***, ventromedial hy. n.; ***VTA***, ventral tegmental area; ***xscp***, crossing of the superior cerebellar peduncle; ***ZI-H***, zona incerta and lenticular fascicles (H fields).

##### 3.1.3.6 Preoptic Area and Hypothalamus

The preoptic complex (POC; levels 2 and 3) and the hypothalamus (Hy; level 2) contain 3 and 17 ROIs, respectively (Table S1). At level 4, the POC bifurcates into the preoptic region (POR) and the subjacent segments of the optic nerve and chiasma (opt), which, despite being functionally unrelated, were artificially merged because they frequently ‘fuse’ at low MRI resolutions. At levels 5 and 6, POR partitions into the different (poorly distinguishable) nuclei of the preoptic area (PO), *per se*, and the medial lamina terminalis, and its vascular organ (drawn together as LTer-VOLT). LTer-VOLT and opt are readily identifiable throughout G12 and NMT v2, due to their starkly distinct contrast and macroscopic location of LTer-VOLT over opt or at the base of PO (Fig. 9a,b). PO is identified by its canonical location at the level of the optic chiasma and along the anterior part of the third ventricle (3V), as well as by its distinct darker contrast. The latter defines rather sharp boundaries with SDB, anteriorly, and with the anterior hypothalamic nucleus (AH), posteriorly, as indicated by the black arrows in Figure 9b.

Most of the larger subdivisions of Hy are distinguishable due to local variations in signal intensity and/or the presence of specific white matter tracts, such as the mammillothalamic tract (mt). For example, the boundaries between AH, the ventral medial nucleus (VMH), and the arcuate nucleus (Arc) were marked by abrupt changes in contrast, similar to the boundary between PO and AH (see black arrows in Fig. 9b). The perifornical (PeF; not shown), retro-mammillary nucleus (RM) and, more particularly, mammillary nucleus are recognizable by their specific relation to the fornix (f) and mt, which are both identifiable as continuous dorsoventral tracts between the thalamus and hypothalamus (Fig. 9b). The mammillary nucleus of the hypothalamus (MM), which typically surrounds f, is also recognizable due to the bulge (mammillary body) that it forms at the base of the diencephalon (Fig. 9d). The different divisions of the lateral hypothalamic region (LHy; level 5) – that is, the lateral nucleus (LH), peduncular lateral nucleus (PLH), and juxtaparaventricular nucleus (JPLH) – are delineated mainly based on their lighter signal, compared to neighboring ROIs (not shown). The limit between the anterior LH and posterior PLH is set at the level at which the fornix reaches the hypothalamus (not shown; see RMBSC4 Fig. 55). The paraventricular nucleus (Pa) and the posterior hypothalamic nucleus (PH) are distinctively darker and located medially, along 3V. The subthalamic nucleus (STh; not shown; abbreviated elsewhere as STN), which is also part of the hypothalamus (Puelles et al., 2013), is consistently identifiable due to a light circular signal located in-between the darker zona incerta (ZI-H; see Section 3.1.3.7) and substantia nigra (SN; see Section 3.1.3.8). Finally, the pituitary (Pit; or hypophysis) is connected to Hy via the distinct medial eminence (MEm), and contains an anterior (APit; adenohypophysis) and a posterior (PPit; neurohypophysis) division, which are both recognizable in the NMT v2 due to much brighter contrast for PPit, compared to APit (Fig. 9c).

The hierarchical grouping of Hy is based mostly on a classical neuroanatomical grouping (Saper, 2012), rather than on ontological grouping (Puelles et al., 2013), due to the non-contiguity of the alar and basal hypothalamic nuclei in the adult Hy. At level 3, Hy is divided into tuberal (THy), posterior (PHy) and pituitary (Pit) groups (Table S1). The tuberal hypothalamus contains the paraventricular hypothalamus (Pa; also singled out as medial tuberal hypothalamus at level 4), as well as the supraoptic hypothalamus (SOpt), the ventromedial hypothalamic nucleus (VMH), the medial eminence (MEm) and the arcuate nucleus (Arc), grouped together as ventral tuberal hypothalamus at level 4, and, finally, the anterior hypothalamic area (AH), the dorso-medial hypothalamic nucleus (DM), and the three distinct lateral hypothalamic nuclei (LH, JPLH, and PLH), grouped together as dorsal tuberal hypothalamus at level 4. The level 3 posterior hypothalamus contains the posterior nucleus *per se* (PH), as well as the prefonical hypothalamus (PeF), the mammillary hypothalamus (MM) and the retro-mammillary hypothalamus (RMM) grouped together as ventral posterior hypothalamus at level 3. Pit forms a separate group at levels 3 and 4, with APit and PPit being considered separately at levels 5 and 6.

##### 3.1.3.7 Epithalamus, thalamus, and prethalamus

The epithalamus (EpiThal; levels 2-3) contains the pineal gland (Pi; levels 4-6) and the habenula (Hb; levels 4-6). Pi forms a distinct round structure at the midline, above the superior colliculus (SC) (not shown). Hb is located posterior and medial to the thalamus. It is recognizable in NMT v2 by its bright and heterogeneous signal (Fig. 9). The thalamus (Thal) contains 34 ROIs at level 6 (Table S1). These ROIs remain listed individually at level 5, except for the anterior thalamus (A) and the capsule of the anterior nucleus (cap) (Fig. 9a), which are then grouped to form the anterior thalamus region (AR) ROI. At levels 4 and 3, the ROIs are grouped into 12 and 6 larger groups, respectively. At level 4, most of the ROIs are grouped based on their connections (e.g., spinal, cerebellar, and palladio-nigral groups) and classical functional attributions (e.g., “non-specific” intralaminar and midline groups). The dorsal lateral geniculate (DLG; abbreviated elsewhere as LGN) and medial geniculate (MG) nuclei remain ungrouped at level 4, due to their size, anatomical distinctiveness, and functional specificity. At level 3, most of the level 4 ROIs are grouped into yet larger entities based purely on their location within Thal. MG and DLG are grouped into a geniculate ROI (GThal). The reticular thalamus (Rt) remains ungrouped until level 2 (Thal) due to its anatomical distinctiveness.

Most level 6 ROIs of the thalamus are distinguishable in G12 and NMT v2. For example, Figure 10 illustrates the delineation and signal contrast of several distinct thalamic ROIs in one coronal slice of NMT v2. The boundary between some ROIs, such as the posterodorsal and posteroventral parts of the ventrolateral nucleus (VLPD and VLPV), had to be based on their theoretical location and topological relationships. But, in most cases, there was a consistent shift in contrast at ROI borders, such as at the boundary between VLPV (darker) and the medial and lateral parts of the ventral nucleus (VPM-VPL; see black arrows in the left side of Fig. 10a). The brighter signal of the VPM-VPL ROI is consistent throughout its anteroposterior extent.

The delineations of the inter-mediodorsal nucleus (IMD), mediodorsal nucleus (MD) and centrolateral and paracentral nuclei (CL-PC) ROIs were marked by rather sharp changes in signal, from darker in IMD to much lighter in CL-PC. The brighter signal in the medial part of MD likely corresponds to its medial portion (MDM in RMBSC4), which could be added in a future version of SARM. Dorsal to IMD, the stria medullaris tract, paraventricular and paratenial nuclei were grouped into one ROI (PT-PV-sm) due to their small individual sizes. Ventral to CL-PC, the centromedial and parafascicular complex (CMn-PF) is identifiable by its darker contrast, which reveals the typical wing-shaped form of CMn-PF, and by the passage of the fasciculus retroflexus (fr). fr is located just ventral to CMn-PF in Figure 10a, but it can be seen crossing CMn-PF in the parasagittal view of NMT v2 in Figure 9a,b and in the coronal view of Figure 11a.

**Figure 11.**
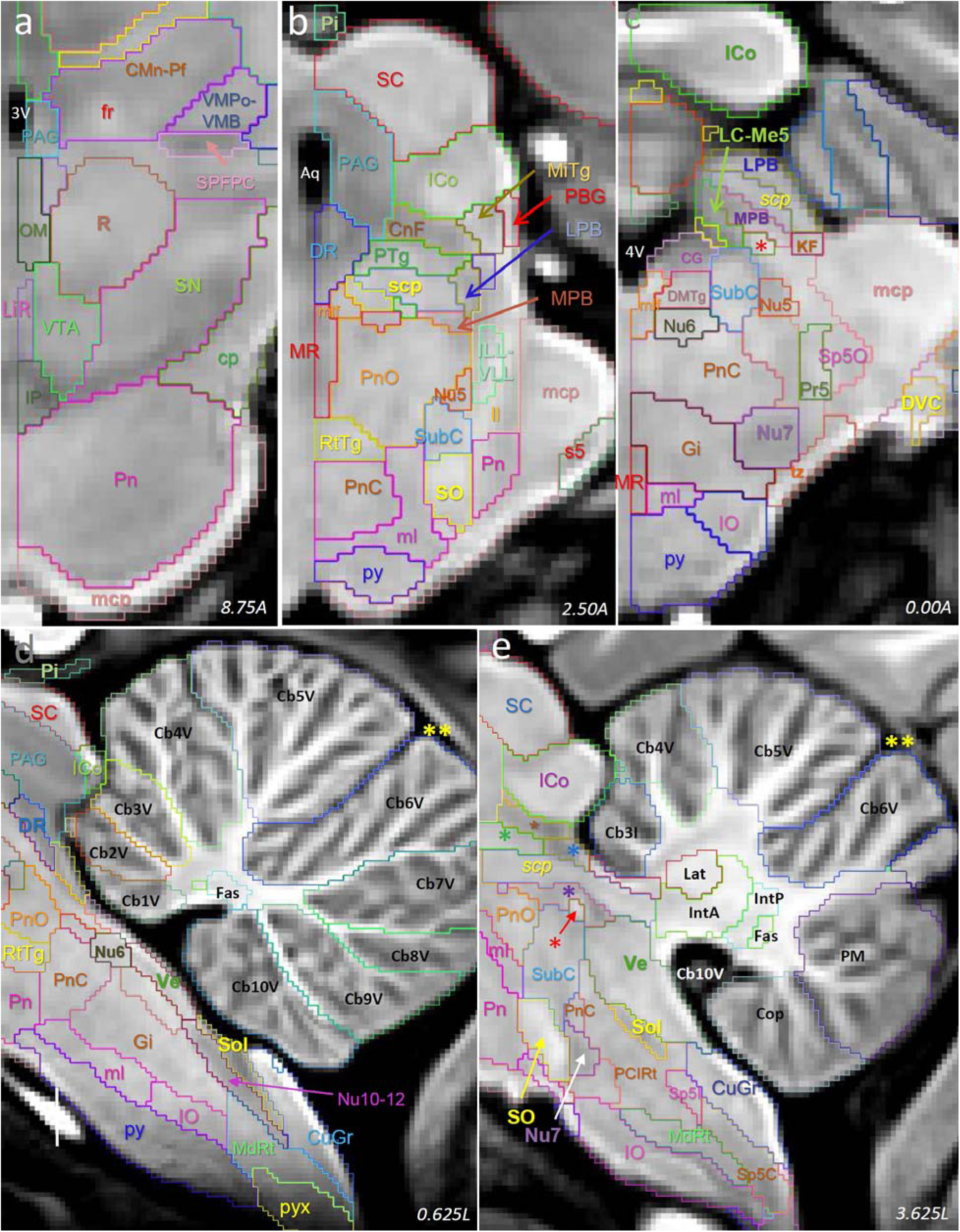
SARM hindbrain ROIs in coronal and parasagittal views. Coronal (**a**-**c**) and parasagittal (**d**,**e**) slices through the NMT v2 showing ROI delineations at various levels of the neuraxis. In (**d,e**), the double yellow asterisks indicate the position of the anterior cerebellar fissure that separates anterior and posterior lobes. In (**e**), the asterisks indicate the locations of CnF (brown), PTg (green), LPB (blue), and MPB (purple). The red asterisk indicates the location of the me5. In (**a-c**), left is medial and top is dorsal. In (**d,e**) left is rostral and top is dorsal. ***Abbreviations***: ***3V***, third ventricle; ***4V***, fourth ventricle; ***Aq***, aqueduct; ***Cb3I***, intermediate part of the cerebellar lobule 3; ***Cb1-10V***, vermis part of cerebellar lobules 1-10; ***CG***, central gray n.; ***CMn-PF***, centromedial and parafascicular thal. n.; ***CnF***, cuneiform n.; ***Cop***, cerebellar copula; ***cp***, cerebral peduncle; ***CuGr***, cuneate and gracile n.; ***DMTg***, dorsomedial tegmentum; ***DR***, dorsal Raphe; ***DVC***, dorsal and ventral cochlear n.; ***Fas***, fastigial (medial) n.; ***fr***, fasciculus retroflexus; ***Gi***, gigantocellular reticular n.; ***ICo***, inferior colliculus; ***IntA***, anterior interposed n.; ***IntP***, posterior interposed n.; ***IO***, inferior olive; ***KF***, Kolliker-Fuse n.; ***Lat***, lateral (dentate) n.; ***LC-Me5***: locus coeruleus and mesencephalic 5 region; ***LiR***, linear Raphe; ***ll***, lateral lemniscus; ***LPB***, lateral parabrachial n.; ***mcp***, medial cerebellar peduncle; ***MdRt***, medullary reticular formation; ***me5***, motor trigeminal root; ***MiTg***, microcellular tegmental n.; ***ml***, medial lemniscus; ***mlf***, medial longitudinal fascicle; ***MPB***, medial parabrachial n.; ***MR***, medial Raphe; ***Nu5***, trigeminal motor n.; ***Nu6***, abducens n.; ***Nu7***, facial n.; ***Nu10-12***, hypoglossal and motor vagus n.; ***OM***, oculomotor complex; ***PAG***, periaqueductal gray; ***PBG***, parabigeminal n.; ***PCIRt***, parvicellular and intermediate reticular n.; ***Pi***, pineal gland; ***PM***, paramedian cerebellar lobule; ***Pn***, pontine n.; ***PnC***, caudal pontine reticulum; ***PnO***, oral pontine reticulum; ***Pr5***, principal trigeminal sensory nucleus; ***PTg***, pedunculopontine tegmentum; ***py***, pyramidal tract; ***pyx***, pyramidal tract decussation; ***R***, red n.; ***RtTg***, reticulotegmental formation; ***s5***, sensory root of the trigeminal nerve; ***SC***, superior colliculus; ***scp***, superior cerebellar peduncle; ***SN***, substantia nigra; ***Sol***, solitary tract n.; ***Sp5C***, caudal spinal trigeminal n.; ***Sp5I***, intermediate spinal trigeminal n.; ***Sp5O***, oral spinal trigeminal nucleus; ***SPFPC***, subparafascicular parvocellular thal. n.; ***SO***, superior olive; ***SubC***, subcoeruleus; ***tz, trapezoid bundle region***; ***Ve***, vestibular n.; ***VMPo-VMB***, posterior and basal parts of the ventromedial thal. N.; ***VTA***, ventral tegmental area.

Ventral to CMn-PF, the posterior and basal parts of the ventromedial nucleus (VMPo-VMb) are identified together as a small, brighter region tucked between CMn-PF and the consistently darker and sharply delimited subparafascicular parvocellular ROI (SPFPC; see also Fig. 9a, 10, and 11a). The ventroposterior inferior nucleus (VPI) ROI, which appears lighter in G12 (not shown), is delineated in the NMT v2 mainly based on its theoretical location at the lateral and ventral base of the thalamus, dorsal to the darker ROI of the zona incerta and H fields (ZI-H). ZI-H, which is the only ROI of the pre-thalamus (PreThal), is characterized by a thin strip of darker signal (see green asterisk in the left side of Fig. 10a), sandwiched at more anterior levels between two lighter strips, likely corresponding to the H1 and H2 fields of the lenticular fascicle (not present at the AP level shown in Fig. 10). The reticular thalamus ROI (Rt) is defined by a thin lighter ‘band’ (in coronal slices) covering the lateral aspect of Thal throughout its rostrocaudal extent.

##### 3.1.3.8 Pretectum and Midbrain

The small pretectum (PrT) and vast midbrain (Mid) contain 4 and 27 ROIs, respectively. The posterior commissure (pc) of the pretectum appears distinctly in the sagittal slice in Fig. 9a,b. The other 3 ROIs of the pretectum are delineated mainly based on their theoretical location in the vicinity of pc, with, however, a slight contrast differentiation for the precommissural nucleus (PrC, Fig. 9a,b) and, to a lesser extent, the posterior commissural nuclei (PCom-MCPC). The midbrain contains several large and distinct ROIs, including the periaqueductal gray (PAG), superior colliculus (SC), inferior colliculus (ICo), and substantia nigra (SN), visible in Figures 9a and 11 (PAG, SC, and ICo). Some smaller ROIs could be delineated based on their distinctively darker or lighter contrast (e.g., interpeduncular nucleus, IP; pedunculopontine tegmentum, PTg; caudal pontine reticulum, PnC; dorsal and median Raphe, DR and MR; superior cerebellar peduncle, scp). Finally, other midbrain ROIs were drawn based on the localization of the aforementioned distinct ROIs. For example, a ventral tegmental ROI was drawn at the base of the midbrain, near its junction with the retro-mammillary nucleus of the hypothalamus (RM), dorsal to the distinctly darker IP, and in between the ventral halves of SN. Lastly, the red nucleus (R) was drawn based on the occurrence of a slight contrast variation forming an ovoid region, dorsal to VTA (Fig.11a).

In the pretectum, pc and two small adjacent ROIs PrC and PCom-MCPC are grouped at level 5 as the posterior commissural region (PCR), to which the prosomeric 1 reticular formation (p1Rt) is added at levels 2-4, to form the PrT ROI. In the midbrain, most level 6 ROIs remain the same at level 5, except for the saginum nucleus ROI (Sag-RL), which joins ICo to form the inferior colliculus complex (ICoC). At level 4, ROIs are grouped mainly based on coarse anatomical or cytological relatedness. For example, SC and ICoC are grouped into a colliculi (Co) ROI; the several tegmental nuclei (e.g., microcellular tegmentum, MiTg, and anterior tegmental nucleus, ATg) are grouped into a midbrain tegmentum ROI (TgMid). Similarly, the midbrain dopaminergic complex (DA-Mid) was formed from the large midbrain dopaminergic cell group ROIs (i.e., VTA, SN and RF) and surrounding structures (e.g., IP). At level 3, these ROIs are further grouped, mainly based on their cardinal location (i.e., dorsal, lateral, medial, and ventral).

##### 3.1.3.9 Pons

The ‘pons’ region of the metencephalon contains 24 ROIs (Table S1), mostly illustrated in Figure 11. The most prominent pons ROI is the pontine nucleus (Pn), located ventrally and well demarcated from the medial cerebellar peduncle (mcp) (Fig. 11a,c,d). The superior olive (SO) forms a bright column posterior to Pn and directly anterior to the darker Nu7 (Fig. 11b, c and e). The lateral and medial parabrachial nuclei (LPB and MPB) form darker bands around the superior cerebellar peduncle (scp), with LPB being posterior to the lighter PTg (Fig. 11b,c,e). Lateral to MPB, we ascribed a small region to a ROI putatively containing both the locus coeruleus and the mesencephalic trigeminal nucleus (Me5). The localization of this ROI is supported by the position of the central gray nucleus (CG), recognizable medial to MPB, and the presence of a small lighter region that corresponds most likely to the efferent trigeminal mesencephalic nerve (me5; marked by the red asterisks in Fig. 11c,e). The lateral lemniscus complex (ll+), which carries projections from the cochlear nucleus, was identified as a light bundle in the lateral portion of the pons, between SO and ICo, which both receive cochlear inputs. Within the boundaries of ll+, we drew ROIs most likely to correspond to the position of the dorsal (DLL) and inferior and ventral (ILL-VLL) lateral lemniscal nuclei (Fig. 11b). Other pons regions, such as the oral and caudal pontine reticulum (PnO and PnC, Fig. 11b-e), were drawn based on their relative position to the structures that were readily identifiable.

##### 3.1.3.10 Cerebellum

The cerebellum (level 2) contains 27 ROIs (Table S1), including 21 cortical ROIs, 4 deep nuclei ROIs, and 2 fiber tracts. The 21 cortical areas consist of the 10 lobules, which are partitioned into the more medial vermis lobules (CbV1 to CbV10; Fig. 11d) and four intermediate lobules (Cb3I to Cb6I; Cb3I is shown in Fig. 11e). The intermediate lobules are continuous with and lateral to the corresponding vermis lobules. The other cortical ROIs are the paramedian lobule (PM), simple lobule (Sim), copula of the pyramis (Cop), ansiform lobules crus 1 (Crus1) and crus 2 (Crus2) ROIs, as well as the floculus (Fl) and paraflocculus (PFl). See Figure 11e for PM and Cop. The deep nuclei are the classical lateral or dentate (Lat), anterior interposed (IntA), posterior interposed (IntP), and medial or fastigial (Fas) cerebellar nuclei (Fig. 11e). While the deep nuclei are clearly revealed by an abruptly darker contrast in G12 (not shown), they are identifiable only by a slightly lighter contrast in NMT v2. This slight increase in lightness is, however, sufficient to delineate the edges of each nucleus. The two tracts are the inferior cerebellar peduncle and olivocerebellar tracts, which are both located close enough to the cerebellum to be allocated to this region, instead of others (unlike mcp and scp, which are mostly represented outside the cerebellum).

At level 5, the vermis lobule ROIs are grouped into anterior and posterior vermis ROIs (AVCbCx and PVCbCx) based on the boundary defined by the primary fissure (double yellow asterisks in Fig. 11d,e) between Cb5V and Cb6V. In addition, the intermediate lobule ROIs, along with Cop, Sim and PM, are grouped into an intermediate cerebellar cortex ROI (ICbCx). Also at level 5, Crus 1 and 2 fuse into a lateral cerebellar cortex (LCbCx) ROI, and Fl and PFl fuse into the Fl-PFl ROI. At level 4, the vermis cerebellar cortex (VCbCx) ROI combines the anterior and posterior vermis ROIs, the deep cerebellar nuclei (DCb) ROI merges the deep nuclei into one, and a cerebellar ‘white matter’ (wmCb) ROI captures the two fiber tracts. At level 3, all the cortical ROIs are grouped under a cerebellar cortex ROI (CbCx), which then coexists with the DCb and wmCb ROIs.

##### 3.1.3.11 Medulla

The myelencephalon (level 1) or medulla (level 2) contains 26 ROIs (Supplementary Table 1). Figure 11d,e and 12 illustrate several of these ROIs. The most obvious ROIs were the solitary tract nucleus (Sol), hypoglossal and motor vagus nuclei (Nu10-12) lying directly ventral to Sol, and the facial motor nucleus (Nu7), due to their sharply delimited darker contrast. The dark Nu7 markedly contrasted against the bright contrast of SO, which lies just anterior to Nu7 (Fig. 11e). The vestibular (Ve) and cuneate-gracile nuclei (CuGr) formed characteristic domes rostral and caudal to Sol, respectively (Fig. 11d). Ventrally, the pyramidal tract (py), decussation of the pyramidal tract (pyx), and inferior olive (IO) were identifiable by their bulging morphology and heterogeneous contrast (Fig. 11d,e; Fig. 12a,b). The cochlear nuclei (DVC, Fig. 12) formed a distinct structure located lateral to the medulla, within the vestibulocochlear nerve (n8). The oral, intermediate (Fig. 12), and caudal spinal trigeminal nuclei (Sp5O, Sp5I, and Sp5C) form a continuous rostrocaudal column made of a medial cellular region (the nucleus itself) and of a lateral fibrous region (the nerve, sp5). Other structures, such as, for example, the paragigantocellular (Gi; Fig. 12) and medullar (MdRt) reticular nuclei, presented a rather homogeneous appearance and were drawn based on their theoretical localization, in between the identifiable regions.

**Figure 12.**
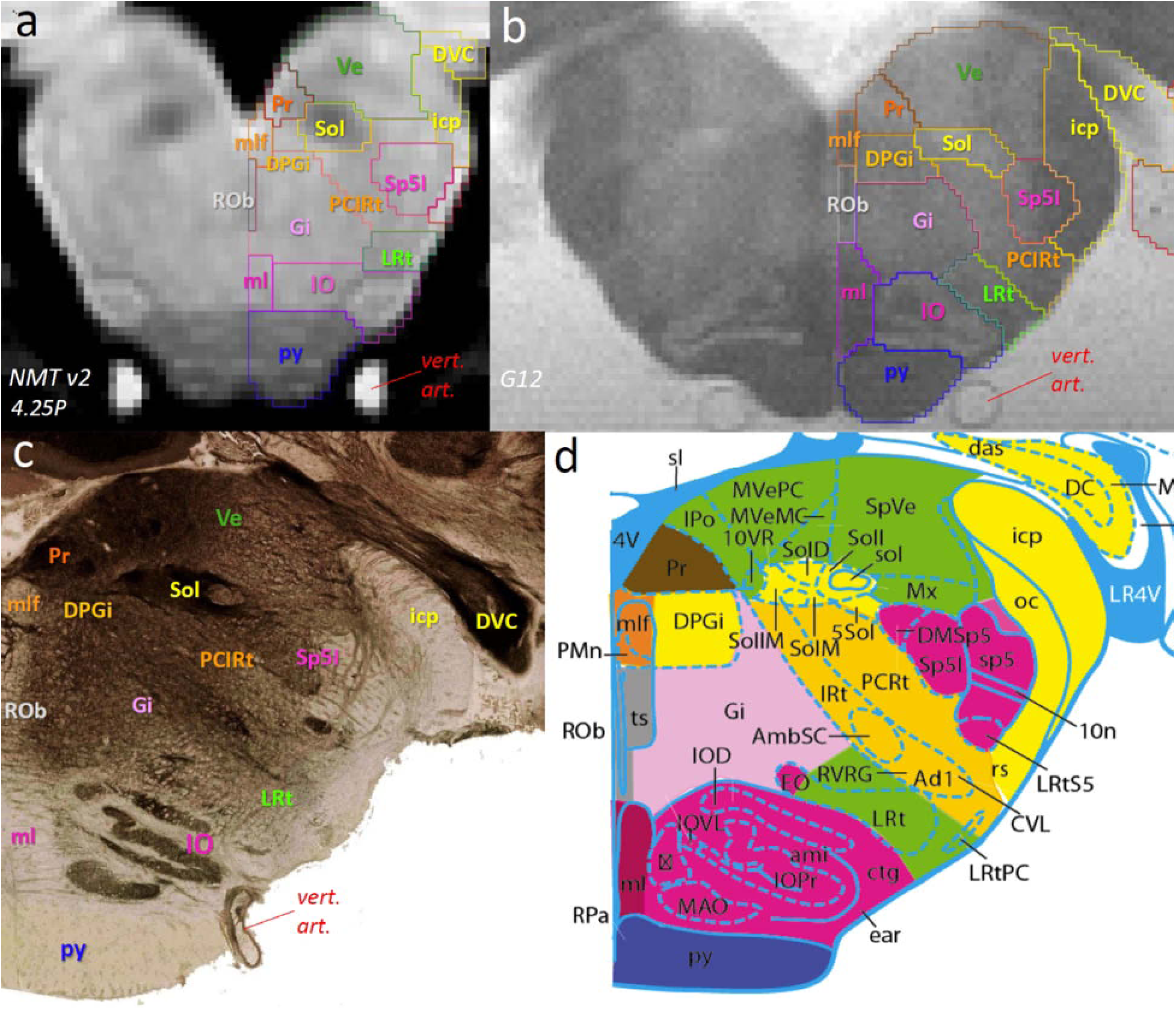
SARM medullar ROIs in coronal view. Coronal slices through the left and right hemispheres of (**a**) the symmetrical NMT v2 and (**b**) the G12. Corresponding RMBSC4 slice through the right hemisphere showing (**c**) an acetylcholinesterase staining and (**d**) its diagram (Fig. 109). ***Abbreviations: DVC***, dorsal and ventral cochlear n.; ***DPGi***, dorsal paragigantocellular nucleus; ***Gi***, gigantocellular reticular n.; ***icp***, inferior cerebellar peduncle; ***IO***, inferior olive; ***LRt***, lateral reticular n.; ***ml***, medial lemniscus; ***mlf***, medial longitudinal fascicle; ***PCIRt***, parvicellular and intermediate reticular n.; ***Pr***, prepositus n.; ***py***, pyramidal tract; ***ROb***, Raphe obscurus n.; ***Sol***, solitary tract n.; ***Sp5I***, intermediate spinal trigeminal n.; ***Ve***, vestibular n. In all panels, left is medial and top is dorsal. For the missing abbreviations in (**d**), see Paxinos et al., 2009, where most abbreviations are similar to Paxinos et al., in preparation.

At level 5, Sp5O, Sp5I, and Sp5C are grouped into a larger spinal trigeminal nucleus ROI (Sp5). At level 4, the ROIs are grouped into 8 composite structures based on their functional relatedness. For example, Ve, DVC, n8, and Pr were grouped in a larger vestibulo-cochlear complex (VCC). Among the 8 composite level 4 ROIs, the medullar Raphe (MedRaphe) and medullar motor nuclei (MedMC) are composed of non-contiguous primary ROIs. Finally, at level 3, the ROIs were grouped based on basic cardinal direction (dorsal, intermediate, and ventral medulla).

### 3.2 Functional Localizer

#### 3.2.1 Individual registration to SARM in NMT v2

The functional localizer data from three rhesus macaque subjects (M1-M3) were nonlinearly registered to the NMT v2 template space for analysis. The quality of the anatomical registration was visually checked (Sections 2.23 & 2.24). As an illustration of the alignment quality, **Supplementary Figure 2** shows the anatomical correspondence between the three macaque subjects and the NMT v2 in the coronal and sagittal planes. Representative coordinates within the periaqueductal gray (PAG, a midbrain region), and the dorsolateral geniculate (DLG) of the right thalamus are labeled.

#### 3.2.1 Subcortical Activation Clusters

A functional paradigm was used as a validation test to determine whether the SARM could sufficiently localize activity to an expected subcortical region. For this, we used fMRI data collected from three monkeys during a visual stimulus paradigm (flickering checkerboard) that was shown by Logothetis and colleagues (1999) to robustly activate the DLG, which is also known as the lateral geniculate nucleus, or LGN. All functional volumes (i.e. time points) collected were included in the analysis. From this analysis (AFNI- and SPM-based), the statistical results were computed across 2 functional scan sessions per individual. For both analysis packages, we found a consistent, bilateral response within and in the vicinity of the DLG (Fig. 13). To compare the extent of functional activity to the anatomically defined DLG, the significant functional activity correlated with the visual flicker stimulus in monkey subject 3 (M3 in Fig. 13a; *p* = 0.05, FDR-corrected) is shown in conjunction with the contour of the DLG in the NMT v2 (SARM levels 4-6; Fig. 13b). In the case of subject M3, almost all of the DLG was activated as determined by the fraction of functionally activated voxels in the atlas DLG (Figure 13c). BOLD activity within this SARM region was consistently positive in all 3 macaques during presentation of the visual stimulus (Figure 13d). The average BOLD percent signal change across the DLG for all subjects and hemispheres was found to be 0.28±0.07% (mean±STD). To illustrate the extent of functional activation spread, a 3D rendering of subject M3’s DLG-localized activation clusters with the atlas DLG (underlaid) was created using SUMA (Saad et al. 2004; Figure 13e). This analysis shows how the SARM can be used for quantifying BOLD activity in individual ROIs and determining the specificity of functional activation. The SARM can, furthermore, be used to assess how an anatomical region responds to a functional manipulation.

**Figure 13.**
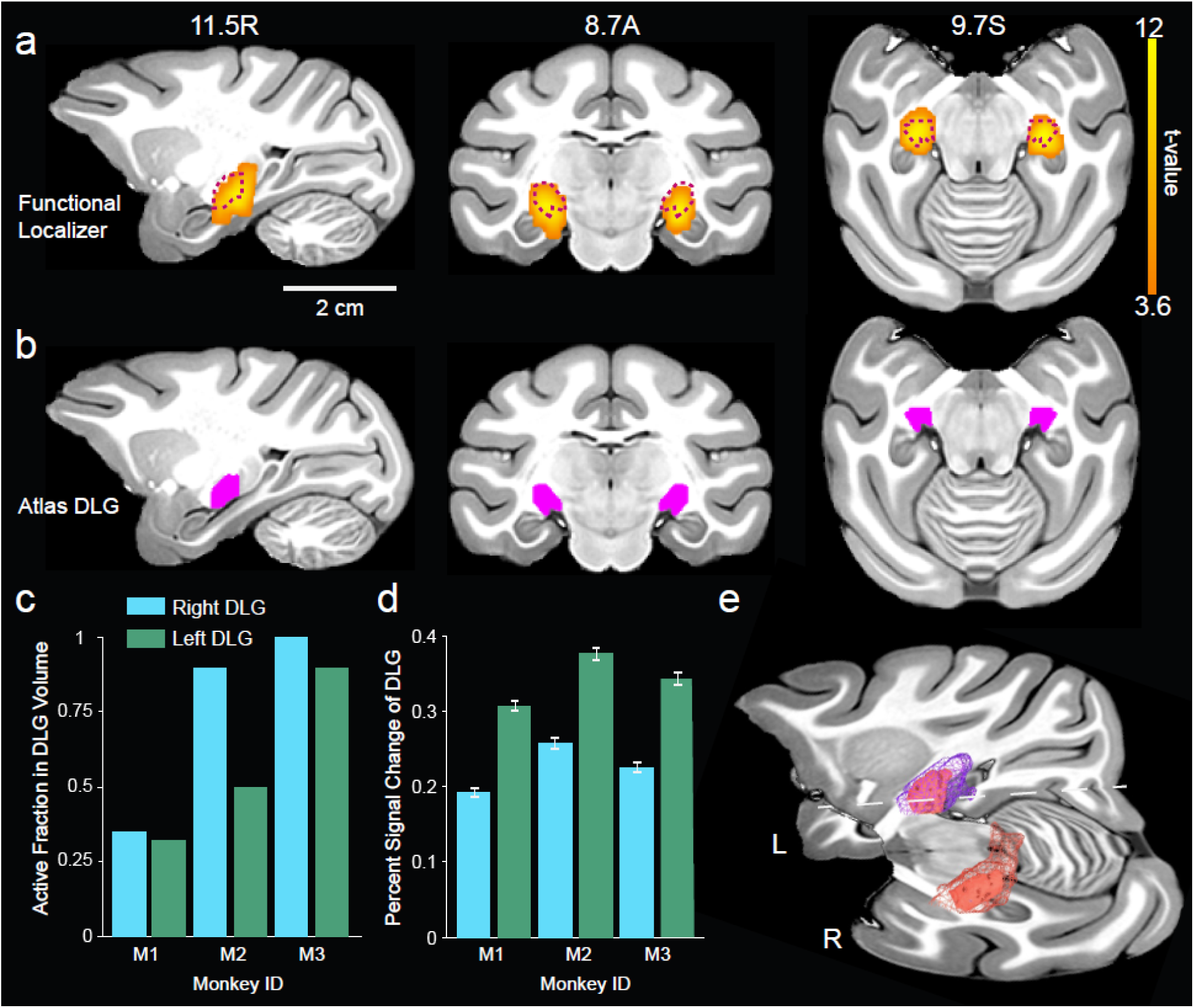
Functional Localizer for DLG. The functional activity elicited in anesthetized monkeys by a flickering checkerboard stimulus was evaluated using the atlas-defined Dorsal Lateral Geniculate (DLG) region. (**a**) Significant positive BOLD activity elicited in monkey M3 is shown on three sections of the NMT v2 volume that include the DLG. Color shows the t-value of significantly activated voxels (FDR-correction at *p* = 0.05; results calculated by the SPM12 analysis pipeline). The anatomical borders of the SARM’ DLG are shown in (**b**) in magenta and with a dashed outline in (**a**). Slice coordinates are in mm relative to the origin (EBZ; ear bar zero). DLG activation in each hemisphere was quantified for 3 macaque monkeys (Monkey IDs: M1-M3) by (**c**) the fraction of functional voxels within the DLG region that were significantly activated (*p* < 0.05, FDR-corrected) and (**d**) the percent signal change (i.e., beta coefficient) associated with the flickering checkerboard (i.e., the 4 sec stimulus ON period) averaged across all functional voxels in the DLG. Error bars plotted represent the standard deviation. (**e**) 3D renderings of the DLG as defined by the atlas (smaller) and functionally by the localizer (larger) displayed in SUMA for monkey M3, against an intersecting axial and sagittal slice (unthresholded, *p* < 0.07; results calculated by the AFNI analysis pipeline).

## 4. Discussion

Here, we have introduced the SARM, a digital neuroanatomical parcellation atlas of the macaque monkey subcortex. This atlas is mapped onto the symmetric NMT v2 population template, which reflects the average morphology of an adult rhesus macaque. The SARM offers a subcortical reference matrix that is suitable for the localization of any neuroimaging results in single-subject and group analyses, as well as for experimental surgical planning. Being in fixed stereotaxic coordinates, the SARM referential remains fixed, regardless of changes in border definitions or the nomenclature of anatomical regions with subsequent optimizations. The atlas was originally drawn on the high-resolution coronal sections of an *ex vivo* MRI of a single subject, with reference to histological material from other subjects, and then manually revised after nonlinear alignment to the *in vivo* population template. Subcortical areas in the forebrain, midbrain, and hindbrain were parcellated according to the Rhesus Monkey Brain in Stereotaxic Coordinates atlas (Paxinos et al., 2009), with revisions that will be reflected in a new edition (Paxinos et al., in preparation). Not all of the small subcortical cytoarchitectonic regions defined in the RMBSC4 (∼900) were drawn. Instead, we incorporated, in larger ROIs, small cytoarchitectonic structures that cannot be identified using the lower MRI resolution, and that would not be pertinent for the localization of BOLD activity. Some of the smaller cytoarchitectonic regions within a single ROI may be functionally unrelated, as they were grouped ‘around’ a larger region, mainly based on their spatial proximity. However, Table S1 lists all the small cytoarchitectonic regions assumed to be included within each SARM ROI. This will allow users to evaluate, based on their paradigm, whether the main or a smaller region might be responsible for the observed BOLD signal.

Refinements of the SARM will be released periodically based on user community feedback. In addition, SARM describes the anatomy at 6 different spatial scales, so that it can be used to name and localize small nuclei, mid-size structures suited to describe fMRI activations, and the major developmental divisions of the subcortex. Finally, we tested and validated the SARM using a DLG functional localizer to localize and quantify BOLD activity with respect to different subcortical regions. Further analyses can essentially be computed using the SARM parcellation presented here, whereby functional activity from any atlas label can be assessed both in terms of areal specificity and sensitivity.

The utility of an MRI atlas is largely determined by how well data can be warped between the native individual space and the common space of the atlas. Data is commonly warped to the common space of an atlas for analysis but may also be warped from the common space to an individual scan. Achieving an accurate registration between the source and target datasets is critical for these processes. By providing the SARM on the *in vivo* population NMT v2, we hope to facilitate simple and accurate alignment between *in vivo* functional data and a template that closely matches its morphology. To obtain an accurate subcortical atlas, special attention was paid here to ROI positioning after alignment of the G12 parcellation to the NMT v2. We assessed this alignment visually on the structural template and examined various interpolation and regularization schemes to minimize errors. Residual inconsistencies between the subcortical labeling and the NMT v2 structure were manually corrected, ensuring an accurate representation of the subcortex in the NMT v2 space.

As there are many spatial scales by which the macaque subcortex can be subdivided, one strategy is to parcellate different brain areas as finely as afforded by cytoarchitectonics. However, for MRI studies, such a fine parcellation is often unnecessary as the discernible differences between structures are limited by image resolution. Further, the fine scale of some cytoarchitectonic structures can be problematic. Small or thin structures are susceptible to large distortions during nonlinear alignment or resampling and may introduce discontinuities or abnormal shapes or cause a region to disappear entirely. Additionally, after resampling to an fMRI grid, some small regions might only consist of a few voxels, making averaging over such ROIs limited, statistically underpowered, and sensitive to registration errors. To avoid this, individual researchers may combine regions to ensure they are robust and adequately sample the desired area, but this can introduce ambiguity. For instance, one researcher’s definition of the regions comprising the amygdala may differ from another, and replicability suffers when researchers lack a consistent definition of brain structures. Atlases circumvent this issue by providing independent, structurally defined regions for ROI analysis and quantification. This capability additionally avoids the potential pitfall of circular analysis, where functional activity is localized to a group of structures, and those structures are then analyzed using the same data (Kriegeskorte et al., 2010).

The SARM addresses both the issues of ROI size and consistency by introducing hierarchical groupings. The fine-to-gross classification provides the specificity needed for use with histological material, high-resolution structural scans and targeted brain interventions as well as larger well-defined composite regions suited for reliable fMRI sampling. The SARM’s finer levels (levels 5 and 6) may have some utility at typical fMRI resolutions but are rather advantageous for detailed structure analysis (e.g., MRI voxel intensity, comparisons to histological material, describing surgical, tracer or pharmacological injection sites). The composite regions of levels 1-4 are sufficiently large to limit the impact of nonlinear registration errors and to include a sufficient number of voxels for averaging over a ROI.

The SARM’s composite ROIs are additionally useful for meta-analyses and cross-species comparisons, as finer parcellations differ between anatomists, species (NHP and human), and individuals. By providing composite structures based on cytoarchitectonics and developmental regions, we defined regions that can be used to analyze a target structure at resolutions suited for structural MRI, diffusion MRI and fMRI studies. Conveniently, the SARM can be used in conjunction with the Cortical Hierarchical Atlas of the Rhesus Macaque (CHARM; Jung et al., this issue). This provides researchers with an additional degree of freedom to study subcortical-cortical relationships at varying scales.

There are a few limitations of the current atlas implementation that could be improved in future iterations. First, while the in-plane resolution of the high-resolution scan (G12) was sufficient for definition of subcortical structures, the out-of-plane resolution (1 mm) limited the available information for tracking these structures in the anteroposterior axis. When these structures were warped to scans with higher out-of-plane resolution, manual adjustments were necessary to resolve discontinuities and inaccurate labeling driven by resampling. Isotropic high-resolution imaging in all dimensions facilitates the creation of digital atlases and their generalizability. Secondly, collecting an *in vivo* scan of the G12 subject would have helped to account for disparities between *in vivo* and *ex vivo* preparations and could have acted as an intermediate target when warping between the high-resolution *ex vivo* scan and the *in vivo* NMT v2. Thirdly, *in vivo* and *ex vivo* multimodal neuroimaging would have been helpful in delineating fine boundaries of subcortical structures and improved alignment to scans with differing contrast. It may also permit detecting structures that were not readily identifiable here (e.g., Ce).

Another consideration unique to NHP imaging is the orientation of the brain in the scanner. Humans are typically scanned in the supine position. However, macaques are scanned in various positions. The “sphinx” position is the most common, but being seated in a vertical scanner is also fairly common. Visual inspection of scans collected in the sphinx and seated positions suggest a change in the brainstem’s orientation with respect to the rest of the brain. Affine alignment is unable to correct for such relative differences in brainstem orientation, and depending on the algorithm, even nonlinear alignment tools may be limited, insofar as the brainstem can be adjusted. Further investigation is required to evaluate how well brainstem structures are registered between a stereotaxic template and functional data collected in the vertical seated orientation. Additionally, it is worth noting that *ex vivo* tissue sections may differ from brain imaging due to differences in features (i.e., brainstem orientation, CSF volume, ventricle size, and sulcal position). In particular, coronal sectioning through the brainstem for histological analysis might be perpendicular to the rostrocaudal axis, whereas for an MR scan, coronal sections are generally oriented with respect to the telencephalon.

Users must carefully consider the issue of alignment between their datasets and the atlas in template space. If alignment is not done properly, a mislocation in ROI assignment can occur. The same notion applies when attempting to warp an atlas to individual scans. The SARM can always be further improved by using additional isotropic high-resolution structural scans, by using Positron Emission Tomography (PET) with chemoarchitectonically specific radiolabeled ligands (e.g., Oler et al., 2012) or by applying other functional localizers (e.g., mechanoreceptive stimuli for localizing activity in thalamic nuclei CuGR and VPL-VPM or auditory stimuli to activate DCV and MG). The SARM atlas regions can be further organized by functional modalities and connectivity-based clustering. It is strongly recommended to examine the vicinity of functional activations and evaluate the relevance of BOLD signal overlap with specific ROIs. Indeed, with the lower spatial voxel resolution of functional scans, and as observed in our localizer validation experiment, significant BOLD signal can spread beyond intrinsic structural landmarks. Level 4 of the SARM hierarchy should be rather safe for most fMRI analyses, but the higher the level, the more careful one has to be to check the quality of fMRI registration.

The SARM is intended to support subcortical localization for a host of neuroimaging datasets (i.e., fMRI, PET, or diffusion imaging). While studies using a small number of macaque subjects can rely on directly comparing their signal location to print atlases, the SARM allows for morphing data from multiple subjects to an MRI-based atlas (and vice versa) to conduct group-level analyses using any neuroimaging modalities. The explosion of community data sharing (Milham et al., 2020) and multi-center NHP fMRI projects should, therefore, highly benefit from this resource. Furthermore, the SARM itself was conceived and developed within the context of the PRIMatE-Data Exchange (PRIME-DE) and will greatly benefit from usage-based feedback from the community. In addition to its utility for data analysis and identifying structures, the SARM in stereotaxic space has the potential to aid with surgical planning in studies involving tracer injections, drug injections, lesions, electrophysiology, optogenetics, and electrical stimulation, including deep brain stimulation (e.g., Ewerts et al., 2017). Beyond studies conducted solely in macaques, providing that homology and nomenclature equivalencies can be reliably established, future harmonized versions of the SARM, CHARM and human atlas counterparts could further help in comparing structural and functional organization between macaques and humans at a broad scale (Mantini et al., 2012). In this context, the detail of the SARM may be especially important, for example for DBS, where experimentally exploring spatially distinct neurostimulating sites could help explain variations in clinical results (Ewerts et al., 2017).

## 5. Conclusion

We have presented a new subcortical atlas for the rhesus macaque: the Subcortical Atlas of the Rhesus Macaque (SARM). Based primarily on the high-resolution MRI of a single subject and comparison with histological materials, this atlas provides the most detailed subcortical parcellation to date and is the first specifically applied to the subcortex available in a digital format for use by the MRI and general neuroscience community. We provide a specific use case and working examples of the use of this atlas within two popular fMRI analysis software packages. The SARM is part of a larger push in the NHP neuroimaging community to share data and resources. Information on the SARM and other macaque resources may be found at the <u>PRIME-RE</u> (Messinger et al., this issue).

**Abbreviations**
fMRI: functional Magnetic Resonance Imaging
DLG: Dorsal Lateral Geniculate
NHP: non-human primate
NMT v2: NIMH Macaque Template
RMBSC4: 4^th^ edition of the Rhesus Monkey Brain in Stereotaxic Coordinates
ROIs: regions of interest
SARM: Subcortical Atlas of the Rhesus Macaque
CHARM: Cortical Hierarchy Atlas of the Rhesus Macaque

## CRediT author statement

**Renée Hartig:** conceptualization; resources; methodology; software; formal analysis; visualization; writing – original draft; writing – reviewing & editing

**Daniel Glen:** resources; methodology; software; formal analysis; visualization; data curation; writing – original draft; writing – reviewing & editing

**Benjamin Jung:** resources; methodology; software; formal analysis; visualization; writing – original draft; writing – reviewing & editing

**Nikos K. Logothetis:** resources; methodology; funding acquisition.

**George Paxinos:** resources; methodology; writing – reviewing & editing

**Eduardo Garza-Villarreal:** resources; methodology; visualization; software; formal analysis; writing – original draft; writing – reviewing & editing

**Adam Messinger:** conceptualization; resources; methodology; software; visualization; writing – original draft; writing – reviewing & editing

**Henry C. Evrard:** conceptualization; resources; methodology; visualization; writing – original draft; writing – reviewing & editing

## Acknowledgments

EAGV would like to thank Gabriel A. Devenyi for his feedback and support, as well as the Laboratorio Nacional de Visualización Científica Avanzada (LAVIS) for the use of their computer cluster and the Laboratorio Nacional de Imagenología por Resonancia Magnética (LANIREM). RH and HCE would like to thank Michael Beyerlein and Thomas Steudel for their technical assistance with 7T imaging, and Yusuke Murayama for discussions on the visual flicker paradigm. This work was funded in part by the Max Planck Society and by the Intramural Research Program of the NIMH and NINDS (ZIA MH002918 and ZICMH002888).

## Supplementary Material

### 1. Supplementary Tables

**Table S1: List of all ROIs and their hierarchy.** <u>See .CSV file.</u>

### 2. Supplementary Figures

**Figure S1.**
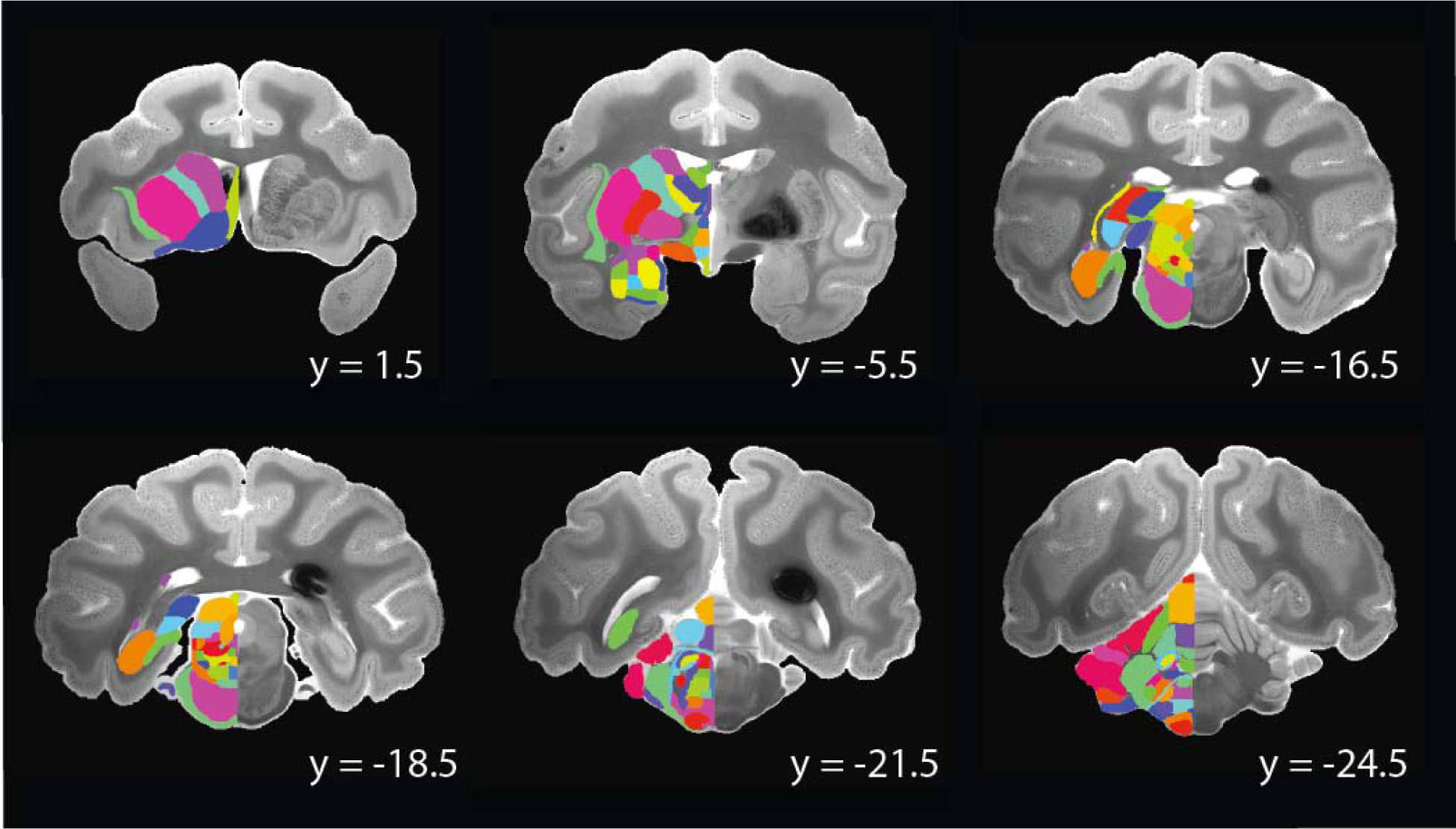
The subcortical atlas in G12 space. Coronal sections from subject G12 are shown with the subcortical parcellation overlaid for the left side regions. The high-resolution *ex vivo* scan was reoriented from its original orientation to a standard orientation. Coordinates are listed in mm in the individual subject’s native space, left hemisphere shown left.

**Figure S2.**
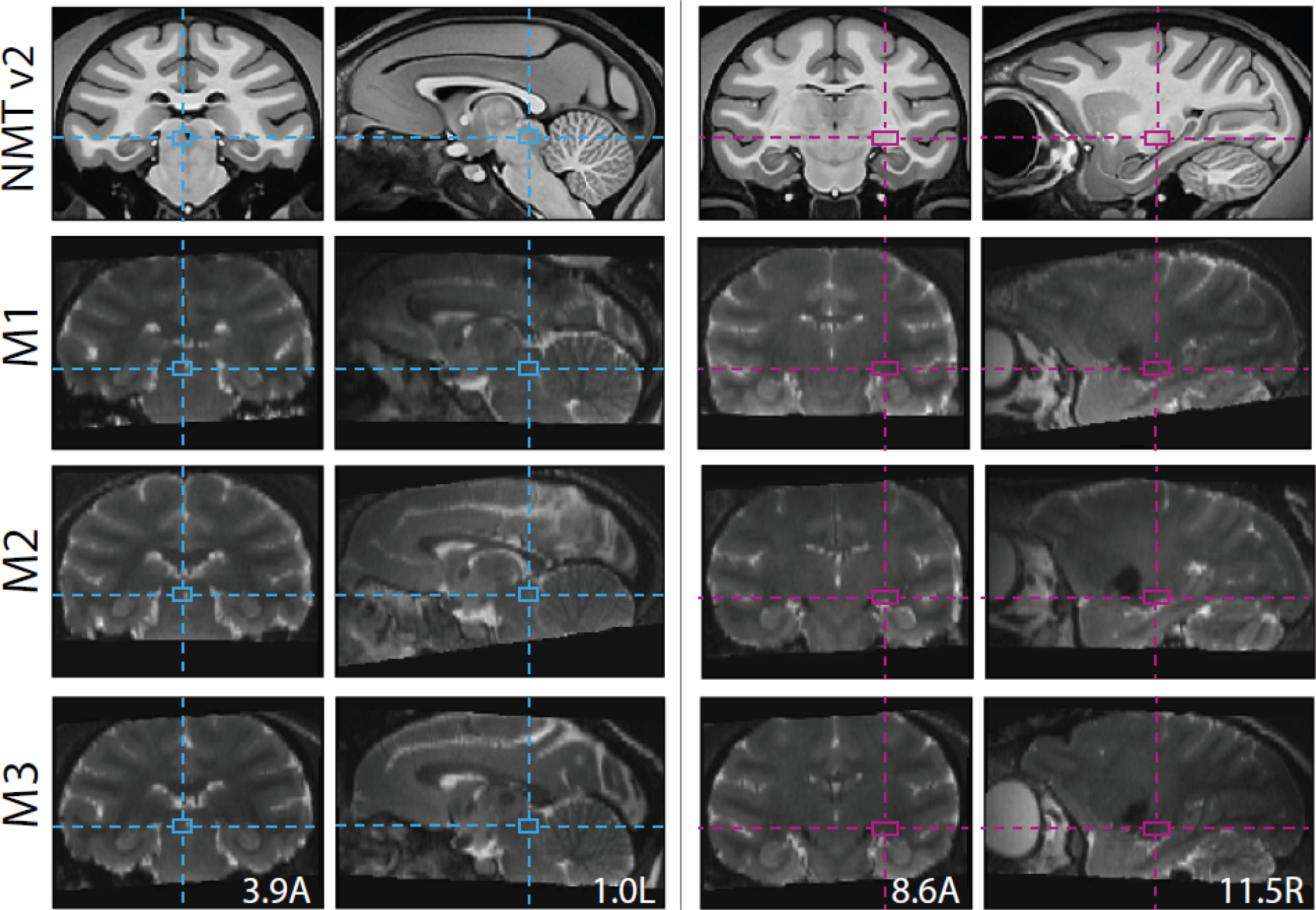
Nonlinear registration of three rhesus macaques to the NMT v2 template. Crosshair intersecting at two subcortical regions: the left periaqueductal gray nucleus (PAG; left panels) in the mesencephalon and the right dorsal lateral geniculate nucleus (DLG, right panels) of the thalamus. The depth of all alignment boxes is 11.6S (stereotaxic coordinates reported in mm from the ear bar zero; EBZ). The coronal and sagittal sections show the correspondence between the T1-weighted NMT v2 and the T2-weighted single-subject anatomical scans from macaque monkeys M1-M3 after Dartels-based nonlinear registration to the population template.

